# Ubiquitination mediates protein localization in RNA virus-infected cells

**DOI:** 10.1101/2024.08.15.608034

**Authors:** Shihua Shi, Vytautas Iesmantavicius, Amit Santhu Sabu, Charlotte Soneson, Hubertus Kohler, Jacint Sanchez, Sucheta Ghosh, Chun Cao, Yong Huang, Gabriele Matthias, Yohei Yamauchi, Patrick Matthias, Longlong Wang

## Abstract

Viruses trigger monocytes’ proinflammatory and antiviral responses. Ubiquitination, a post-translational modification primarily marking proteins for degradation, regulates cellular responses to virus infection. However, a comprehensive analysis of virus-induced ubiquitination in monocytes is lacking. Here we identified a widespread increase of ubiquitination under viral RNA challenge or influenza infection in monocytes. Systematic proteome studies revealed that influenza infection elicits dynamic ubiquitinome alterations, with a notable transition from early to late stage. Most of this increased ubiquitination is not proteolytic and targets proteins involved in subcellular localization, such as the mitochondrial protein COA7 which, when ubiquitinated during infection, translocates to the nucleus and inhibits stress granules formation and TNF-α expression. Blocking ubiquitination halts viral ribonucleoprotein’s nuclear export, highlighting ubiquitination’s importance for protein localization during virus infection.

## Main Text

Ubiquitination is a post-translational modification (PTM) where the small protein ubiquitin (Ub) is attached to a target protein via an isopeptide bond. Depending on the number of added Ub molecules, ubiquitination can be classified into mono-ubiquitination and poly-ubiquitination. Poly-ubiquitination is further categorized by linkage types, in function of the lysine residues that are used for polymerization, such as K29, K48, and K63 among others (reviewed in (*1*)). A major function of ubiquitination is to target misfolded or damaged proteins for degradation via the Ub -proteasome system (UPS). This is achieved by the addition of K48 branched chains to lysine residues of the target proteins which are then degraded by the proteasome (*2, 3*). The proteolytic K48-linked poly-ubiquitination is a major component of the cellular protein quality control (PQC) system. Non-proteolytic ubiquitination like linear ubiquitination, mono-ubiquitination and K11- or K63-linked poly-ubiquitination regulates multiple cellular functions, such as signaling pathways (e.g., immune signaling) (*4, 5*), cell fate decisions (*6*), or programmed cell death (*7*). Both proteolytic and non-proteolytic ubiquitination functions are critical for cellular responses to microbial pathogens (e.g., bacteria and viruses). For example, K63-linked ubiquitination regulates receptor-interacting serine/threonine-protein 1 (RIP1) and RIP2’s interactions with Toll-like receptor (TLR) 3, activating antiviral signaling pathways (*8*) and proinflammatory pathways (*9*). In the inflammatory signaling pathway controlled by the transcription factor nuclear factor (NF)-κB, K63-linked ubiquitination on NF-κB essential modulator (NEMO)’s zinc finger domain mediates its interaction with other molecules and promotes the activation of the pathway (*10*). Additionally, the proteolytic ubiquitination of NF-κB inhibitor, inhibitor of kappa B (IκB), leads to its proteasome dependent degradation, further accelerating the inflammation (*11, 12*). Therefore, understanding the cellular ubiquitination state globally and its functions during pathogen infections is important.

Eukaryotic cells have intricate pathways to respond to exogenous stresses, such as heat-shock, oxidative stress, proteasome stress, or virus infection (*13*). A typical stress response entails translation pausing (*14*) and cellular physiology changes such as RNA splicing inhibition (*15, 16*), ribosome ubiquitination (*17*), stress granules (SGs) formation (*18*), and misfolded proteins management (*19*) among others. Recently, a global increase in protein ubiquitination in epithelial cells was observed under heat-shock (*20*). Blocking this ubiquitination was found to disturb the cellular recovery from stress, concomitant with delayed disassembly of heat-shock induced SGs. Virus infection is a pathogenic stress triggering various cellular responses, which are different for DNA and RNA viruses and show great specificity. For DNA viruses, viral DNA is recognized by cyclic GMP-AMP synthase (cGAS), which further recruits the stimulator of interferon genes (STING) and triggers inflammatory genes expression or activates defense mechanisms (*21, 22*). In the case of RNA viruses, e.g. influenza A virus (IAV) and severe acute respiratory syndrome coronavirus 2 (SARS-CoV-2), different viral genomic sensors, such as TLRs, retinoic acid-inducible gene I (RIG-I)-like receptors, and melanoma differentiation-associated protein 5 (MDA5), are responsible for viral RNA recognition (*23-25*). In the past 5 years, SARS-CoV-2 has been an immense public health threat: until now, over 750 million people have been infected by this coronavirus globally and more than 7 million have died (*26*). Thus, given the importance of ubiquitination for innate immune responses and inflammation, analysis of RNA virus induced global protein ubiquitination can provide important novel insights into proinflammatory responses.

It has been shown that SARS-CoV-2 infection leads to changes on the landscape of ubiquitinated proteins, or “ubiquitinome”, including the ubiquitination of 263 proteins and the deubiquitination of 105 proteins at 24 hours (hr) post-infection in lung epithelial A549 cells (*27*). Notably, SARS-CoV and SARS-CoV-2 induce different ubiquitination patterns, indicating that the virus-induced ubiquitination is virus-specific. Another study of SARS-CoV-2 infection in lung epithelial Calu3 cells revealed that the host cellular ubiquitination system is used by coronavirus to facilitate virus infection by ubiquitinating the viral spike protein (*28*). Although epithelial cells are commonly regarded as the primary targets of respiratory RNA viruses, recent studies have highlighted the susceptibility and importance of monocytes as viral targets and reservoirs, as these cells promote inflammation (*29, 30*) (reviewed in (*31*)). An increase in CD14^high^CD16^+^ intermediate monocytes, positive for SARS-CoV-2 virus, is observed in the blood of COVID-19 patients. Within the infected monocytes, the NLRP3 inflammasome pathway is activated, promoting tissue inflammation (*32*). Similarly, highly pathogenic influenza H5N1 can infect mice or human monocytes and elicit a high level of proinflammatory cytokines in the lung (*33*). In IAV infected patients, there is a notable increase in both classical and intermediate monocytes in the nasopharynx. The isolated patients’ immune cells reveal that monocytes are the main source of TNF-α secretion (*34*). Investigations into IAV infected mice have shown similarly that circulating monocytes are recruited from the bloodstream and accumulate in lungs during the early phase (2-5 days post-infection) (*35, 36*). While monocytes have been recognized to be the principal source of TNF-α during IAV infection (*34*), there has been no ubiquitinome analysis yet of how ubiquitination is modulated in response to virus infections in these cells.

In this study, we employed comprehensive proteomic analyses using various approaches to examine ubiquitination in response to viral RNA and virus infection in monocytes. We identified a dramatic upregulation of global ubiquitination in monocytic THP-1 cells treated with the viral mimic polyinosinic-polycytidylic acid (Poly (I:C)) or infected with IAV. Our proteomic data, collected at multiple timepoints, provide a unique view of the dynamic changes in ubiquitinated proteins, and reveal a dramatic transition of total Ub conjugates from the early to late stages of infection: the global ubiquitination level, identified by the signature peptide (K-ε-GG) abundance, increases in response to viral RNA challenge, and then decreases within 24 hr. Gene ontology (GO) analysis revealed that proteins involved in protein localization processes are highly ubiquitinated. We further explored the proteolytic and non-proteolytic roles of IAV induced ubiquitination by focusing on eukaryotic translation initiation factor 4H (EIF4H) and Cytochrome c oxidase assembly factor 7 (COA7). Interestingly, non-proteolytic ubiquitination of COA7 regulates its cellular localization, which affects inflammatory cytokine TNF-α expression and SGs formation. Finally, by examining the nuclear export of IAV proteins and RNA, we found that the cellular ubiquitination state determines the localization of viral proteins and RNA as well. Taken together, our findings indicate that ubiquitination mediates protein localization in RNA virus-infected cells.

## Result

### Global ubiquitination is upregulated in Poly (I:C) treated monocytes

Viruses exhibit significant diversity, varying in genome type (e.g., DNA vs RNA, single-stranded vs double-stranded, negative-sense vs positive-sense) and particle structure (enveloped vs. non-enveloped)(*37*). Despite these differences, their genomes activate cellular sensors for exogenous DNA or RNA in a similar manner. To investigate virus-induced ubiquitination, we used the Poly (I:C) and Poly (deoxyadenylic-thymidylic) acid sodium salt (Poly (dA:dT)) to simulate viral RNA and DNA respectively. Both mimics were transfected into the human monocytic cell line THP-1 and the human lung epithelial cell line A549 and we assessed the ubiquitination signal by Ub immunoblotting. Remarkably, in THP-1 cells, a substantial increase in ubiquitination signals in the high molecular weight (HMW) region was observed at 6 hr post Poly (I:C) treatment (Fig. 1A). This increase in ubiquitination was reproducible in Poly (I:C) treated mouse primary bone marrow derived macrophages (BMDMs) (Fig. 1B). However, no significant increase in ubiquitination was observed in either Poly (I:C) treated A549 cells or Poly (dA:dT) treated THP-1 cells (fig. S1A and B), indicating that the ubiquitination increase is specific for monocytes in response to viral RNA challenge. To test whether this ubiquitination increase is important to elicit proinflammatory and antiviral responses, we used inhibitors interfering with ubiquitination at different stages of the process: the pan-E1 ligase inhibitor TAK-243, the proteasome inhibitor Bortezomib, and the deubiquitinase (DUB) inhibitor PR-619. We then measured the protein or mRNA level of the proinflammatory cytokine TNF-α (Fig. 1C), the viral RNA sensor RIG-I/DDX58, and the antiviral cytokines interferon-β (IFN-β)/IFNB1 and interferon-induced protein with tetratricopeptide repeat 2 (IFIT2) (fig. S1C-G). The three inhibitors of the ubiquitination process robustly suppressed the expression of all these genes. This demonstrates a critical role of ubiquitination for proinflammatory cytokine release and antiviral response in monocytes.

**Fig. 1.**
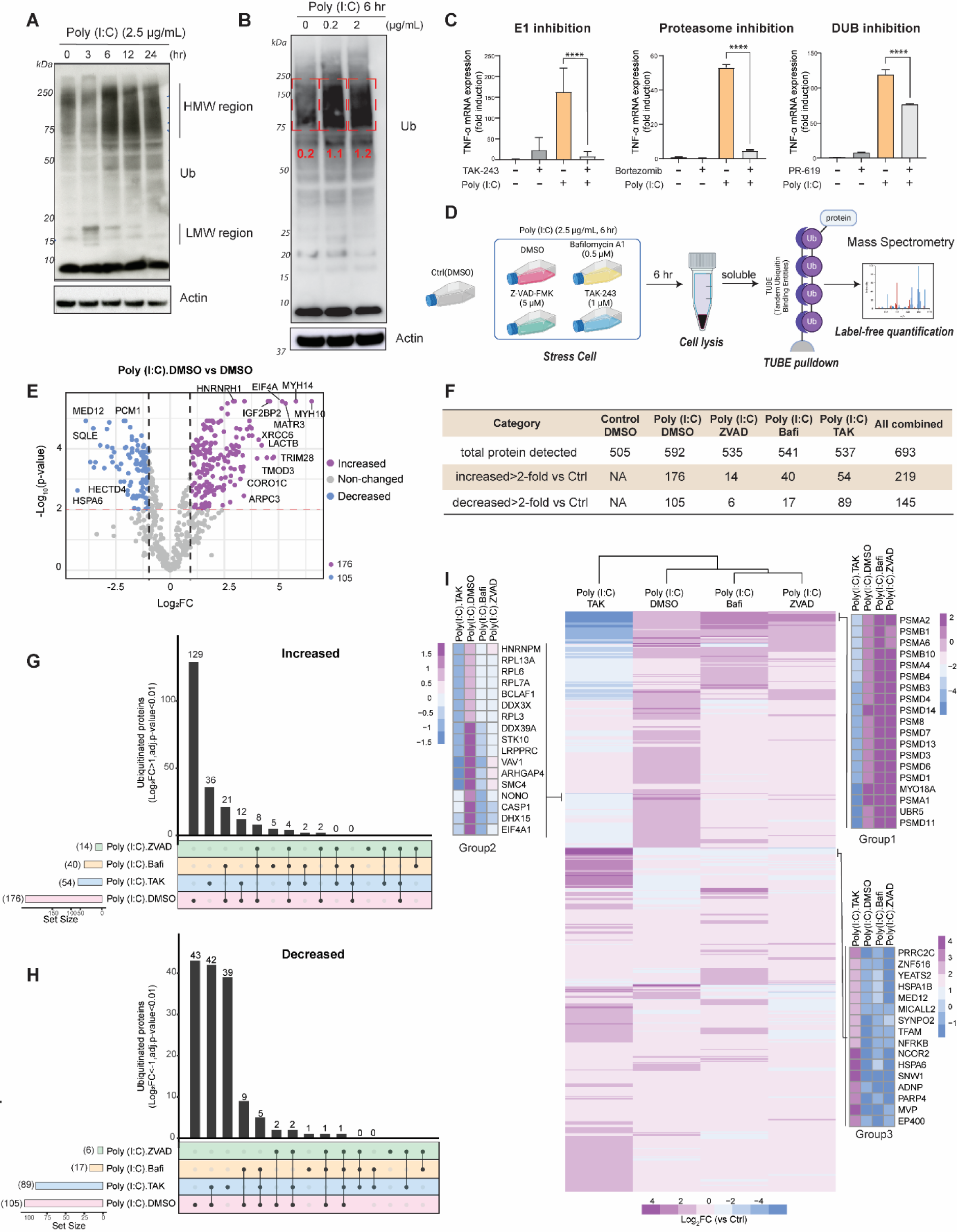
Poly (I:C) induces a global ubiquitination increase in THP-1 cells at 6 hr post-treatment. (**A**) Immunoblotting of global ubiquitination in Poly (I:C) treated THP-1 cells. Poly (I:C) treatment time varied from 0 to 24 hr, as indicated. Actin was used as internal control. (**B**) Immunoblotting of global ubiquitination in Poly (I:C) treated (as indicated) BMDMs. Actin served as loading control. Red numbers are the relative signal intensities in the indicated red dashed rectangles. (**C**) RT-qPCR quantification of TNF-α mRNA level (n = 3). THP-1 cells were treated with 2.5 µg/mL Poly (I:C) together with TAK-243 (1 µM, E1 ligase inhibitor), Bortezomib (1 µM, proteasome inhibitor) or PR-619 (1 µM, deubiquitinases inhibitor) for 6 hr. The 2^-ΔΔct^ method was used for mRNA relative quantification. Statistical analysis was performed with one-way ANOVA test, ****, p < 0.0001. (**D**) Experimental design for the TUBE pulldown-based label-free mass spectrometry analysis in THP-1 cells. Ctrl: Control, only DMSO treated. (**E**) Volcano plot of TUBE based mass spectrometry results in Poly (I:C) treated THP-1 cells. The control group was the TUBE pulldown with DMSO treated cells. Dashed lines represent the log_2_FC = ±1, and p-value = 0.01. Proteins with increased or decreased ubiquitination are displayed based on the threshold log_2_FC > 1 or log_2_FC < -1 respectively (adj.p-value < 0.01), and the numbers of increased and decreased proteins are listed at right bottom corner. (**F**) Summary of all the TUBE mass spectrometry results under different treatments. Proteins with increased or decreased ubiquitination (Log_2_FC > 1 or < -1, adj.p-value < 0.01) were compared to the control group (vehicle treated cells). (**G, H**) Upset plots of proteins with increased and decreased ubiquitination in all the conditions. (**I**) Heatmap of all the proteins (n = 693) Log_2_FC (compared to Control). Enlarged heatmaps are presented for the exemplified proteins in groups 1, 2 and 3.

To obtain a detailed view of the proteins ubiquitinated after Poly (I:C) treatment, we employed Tandem Ubiquitin Binding Entities (TUBE) proteins (*38, 39*) (Fig. 1D and fig. S2A) to selectively capture ubiquitinated proteins from THP-1 cell lysates, followed by label-free quantification with mass spectrometry. In addition to Poly (I:C) treatment, various inhibitors including TAK-243, the pan Caspase inhibitor Z-VAD-FMK, and the endosome maturation inhibitor Bafilomycin A1, were used as multiple controls to investigate the specificity of Poly (I:C)-induced ubiquitination (Fig. 1D and table S1). We detected 500∼600 proteins for each condition, amounting to a total of 693 proteins across all conditions. We examined the proteins with increased (Log_2_Fold-change (FC) > 1, adj.p-value < 0.01) or decreased (Log_2_FC < -1, adj.p-value < 0.01) abundance (indicative of proteins with increased or decreased ubiquitination) relative to control cells (no Poly (I:C) and only vehicle was added). Depending on the condition, 14 to 176 proteins were increased (Fig. 1F). Among these, 176 proteins increased following Poly (I:C) treatment (Fig. 1E, purple dots); this number was reduced to 14 or 40 when Z-VAD-FMK or Bafilomycin A1, respectively, was included in addition to Poly (I:C), suggesting that interfering at different stages of the TLR3 pathway inhibits the ubiquitination (Fig. 1F). In the presence of TAK-243, only 54 proteins were increased, with 36 out of these 54 (67%) being unique as compared to 5 in Bafilomycin and none in Z-VAD-FMK treatment (Fig. 1G). Similarly, Poly (I:C) treatment led to 105 decreased ubiquitinated proteins. Depending on the inhibitor added, this number was reduced to 89 in the presence of TAK-243, or even only 6 in the presence of Z-VAD-FMK (Fig. 1F and H). Based on the principal component analysis (PCA) of all the samples (fig. S2B), it appears that TAK-243 is less specific than Z-VAD-FMK (*40*) and Bafilomycin A1 (*41*) at interfering with Poly (I:C)-activated pathways. Among the 176 increased proteins, 129 proteins were increased only under Poly (I:C) condition (Fig. 1G). These represent the proteins specifically ubiquitinated in response to Poly (I:C).

Unsupervised hierarchical clustering revealed the details of the ubiquitinome variation (compared to control cells) under different inhibitors’ treatments (Fig. 1I). The ubiquitination changes under TAK-243 treatment were different from those under the other conditions (e.g., Group 1 and Group 3 in Fig. 1I). The Group 2 proteins (Fig. 1I), which were selected from the 129 proteins specific to Poly (I:C) treatment, comprised proteins EIF4A1, NONO, and DHX15 among others, which were previously identified in the heat-shock induced ubiquitinome (*20*), as well as antiviral proteins such as CASP1 (*42*) and DDX39A (*43*). GO_Biological Process (BP) (*44*) analysis showed that many of these 129 proteins were connected to RNA-related processes, e.g., translation and mRNA processing (fig. S2C).

### Global ubiquitination varies in a time-dependent manner

To comprehensively explore the global ubiquitination induced by viral RNA, we used tandem mass tag (TMT)-quantitative proteomics to deepen the analysis of the ubiquitinated proteins enriched by TUBE at different timepoints (3 hr, 6 hr, 12 hr, and 24 hr) after Poly (I:C) treatment (Fig. 2A). In comparison to label-free quantification, we successfully identified 4,252 proteins across all the timepoints (table S2). At the earliest timepoint (3 hr), we detected only 8 ubiquitinated proteins with increased abundance (Log_2_FC > 1, adj.p-value < 0.01), and none with decreased abundance (Log_2_FC < -1, adj.p-value < 0.01) (Fig. 2B). This observation is consistent with our immunoblotting data (Fig. 1A) showing no detectable increase in ubiquitination at 3 hr after Poly (I:C) treatment. In contrast, at 6, 12 and 24 hr, our analyses (Fig. 2C to 2E) demonstrated considerable variations in the increased or decreased ubiquitination.

**Fig. 2.**
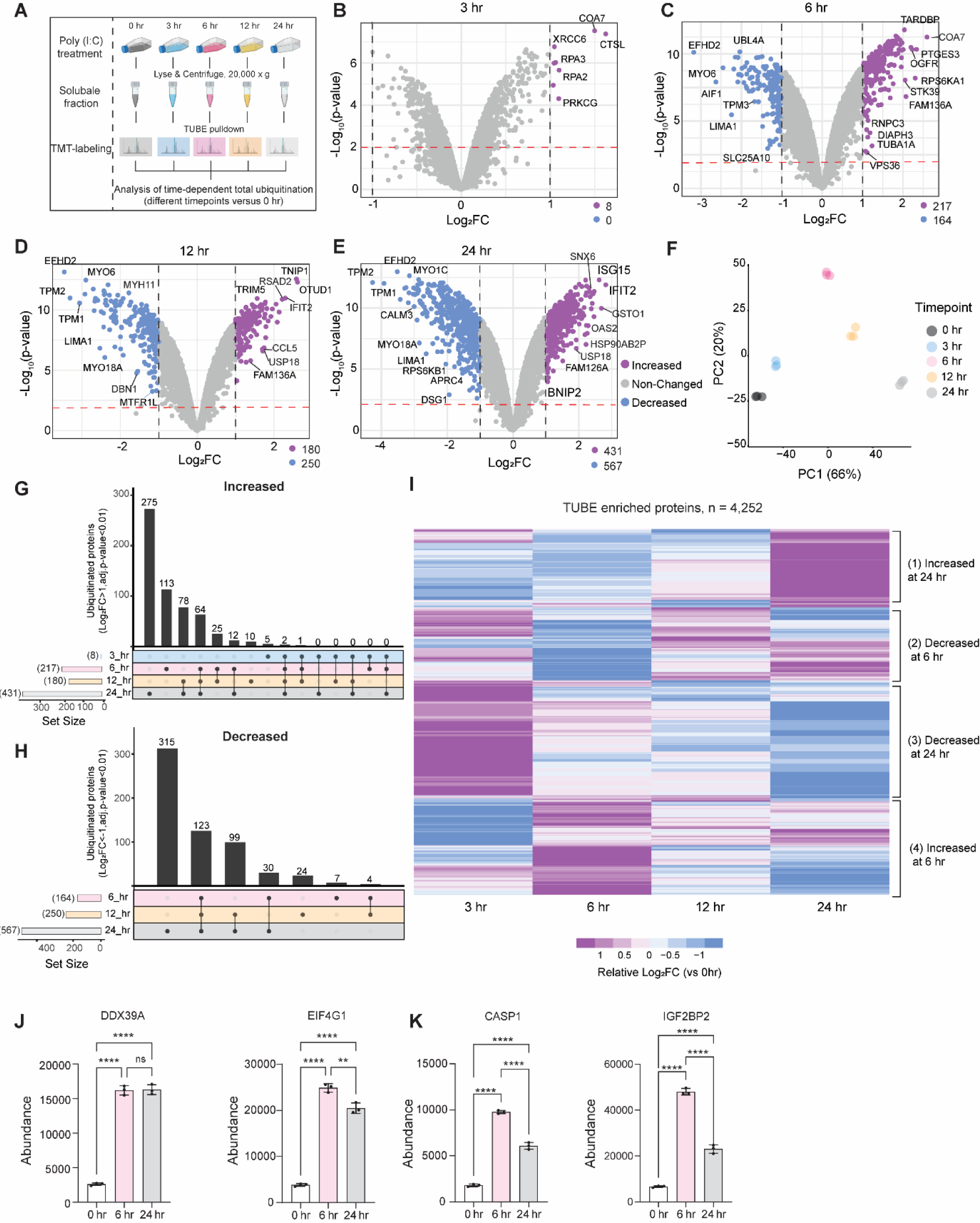
TUBE-TMT reveals distinct ubiquitinomes at different timepoints post Poly (I:C) treatment. (**A**) Workflow for samples from THP-1 cells with different Poly (I:C) treatment (2.5 µg/mL) time. (**B** to **E**) Volcano plots of all the TUBE-TMT analyses at different timepoints after Poly (I:C) treatment. Dashed lines represent the log_2_FC = ±1, and p-value = 0.01. The numbers of proteins with increased and decreased ubiquitination (Log_2_FC > 1 or < -1, adj.p-value < 0.01) are indicated at right bottom corner. (**F**) PCA analysis of all the samples from 5 timepoints: 0 hr, 3 hr, 6 hr, 12 hr and 24 hr. (**G, H**) Upsets plot of proteins with increased (Log_2_FC > 1) and decreased (Log_2_FC < -1) ubiquitination (adj.p-value < 0.01) at all the timepoints. (**I**) Heatmap of all the proteins relative Log_2_FC (compared to 0 hr). The total proteins were categorized into 4 different groups (indicated at right side). (**J, K**) Examples of proteins with increased ubiquitination (based on by the peptide abundance quantified by TMT labeling) at 6 hr post Poly (I:C) treatment. Statistical analysis was performed with one-way ANOVA test, **, p < 0.01; ****, p < 0.0001; ns, not significant.

Across all three timepoints, there were 217, 180, and 431 proteins exhibiting increased ubiquitination (Log_2_FC > 1, adj.p-value < 0.01), while 164, 250, and 567 proteins showed a decrease, as compared to 0 hr (Fig. 2B to E and table S2). In the PCA plot, the ubiquitinomes at 6 and 24 hr were the most distant from 0 hr at PC2 and PC1 respectively (Fig. 2F), which prompted us to investigate these two timepoints further. Next, we determined the number of proteins with increased ubiquitination (Log_2_FC > 1, adj.p-value < 0.01) or decreased ubiquitination (Log_2_FC < -1, adj.p-value < 0.01) at different timepoints. Notably, for proteins with increased ubiquitination, only 78 (64+12+2, 35.8% of 217 and 18.2% of 431) were detected at both 6 hr and 24 hr (Fig. 2G), suggesting a dramatic transition of the Ub conjugated proteins between 6 and 24 hr. In contrast, at 12 hr, except for 10 proteins, all the others were detected at either 6 hr or 24 hr, indicating that 12 hr represents an intermediate timepoint for ubiquitination. The situation was different for decreased ubiquitination (Fig. 2H). From 6 hr to 24 hr, 123 proteins were shared by all the timepoints, accounting for 75%, 49.2%, and 21.7% of proteins with decreased ubiquitination at 6 hr (164), 12 hr (250), and 24 hr (567), respectively. This suggests a continuous deubiquitination process from 6 to 24 hr. Lastly, we validated the selectivity of TUBE pulldown by immunoblotting. For this we chose Myosin 10 and α-tubulin, two abundant proteins which we had identified in our analysis (fig. S3A and B).

To gain a better understanding of the changes in TUBE-identified ubiquitinome, we constructed a heatmap of all timepoints based on Log_2_FC (compared to 0 hr). Comparison of FC patterns at 6 hr and 24 hr showed that they appeared almost reversed (Fig. 2I). Next, based on the variation between these two timepoints, the identified proteins were roughly categorized into four groups: those that (1) increased at 24 hr, (2) decreased at 6 hr, (3) decreased at 24 hr, and (4) increased at 6 hr. Within the group (4) “increased at 6 hr”, we highlighted several proteins which were also identified by label-free quantification (table S1) with varying ubiquitination patterns (Fig. 2J and K). For instance, EIF4G1 and DDX39A exhibit strong enrichment at 6 and 24 hr, whereas CASP1 and IGF2BP2 are highly enriched primarily at the 6 hr timepoint. Further, we examined by GO analysis to which biological processes the 6 hr and 24 hr proteins with increased ubiquitination belonged to. Interestingly, the top 20 processes of these two timepoints were very different with only one process in common (fig. S3C to E). In summary, our findings indicate that the global ubiquitination induced by Poly (I:C) varies in a time-dependent manner, and there is a dramatic transition in proteins with increased ubiquitination between 6 and 24 hr.

### Paired diGly and TMT quantitative proteomics reveal a ubiquitination state transition from 6 to 24 hr

TUBE based affinity purification is efficient to capture poly-ubiquitinated proteins, but some contaminations by Ub chains interacting proteins and loss of mono-ubiquitinated proteins are inevitable. To validate and further deepen our findings from TUBE proteomics, we employed a complementary approach, diGly peptides enrichment. Unlike TUBE based proteomics, diGly proteomics relies on the enrichment of the signature Lys-ε-Gly-Gly (diGly) peptides generated from ubiquitinated proteins or Ub chains following trypsin digestion (*45-47*). To understand the ubiquitination transition between 6 and 24 hr, we monitored the variation of diGly peptides in comparison to the 0 hr timepoint. In parallel, we examined total protein abundance by TMT labeling, to account for potential changes in absolute protein levels that would impact diGly peptides abundance (Fig. 3A). In total, our analysis identified 8,461 proteins, and, as illustrated in Figure 3B and C, we observed merely 11 proteins (0.13% of 8,461) at 6 hr and 106 proteins (1.2% of 8,461) at 24 hr changing significantly in abundance (Log_2_FC > 1 or < -1, adj.p-value < 0.01) (Fig. 3B and C and table S3). This indicates that the abundance of most ubiquitinated proteins remains stable during Poly (I:C) treatment, and that in this setting the induced ubiquitination does not lead to widespread protein degradation.

**Fig. 3.**
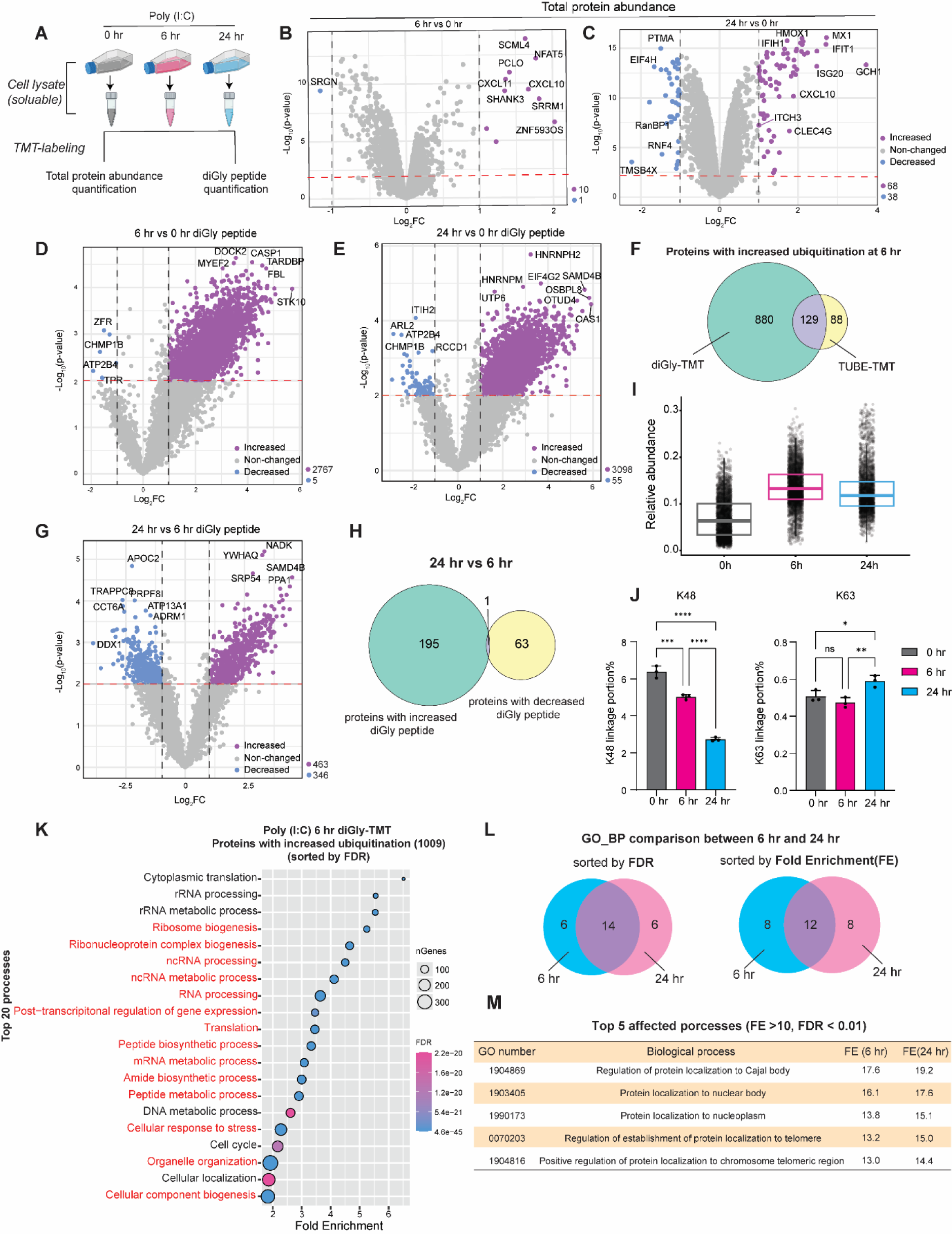
DiGly-TMT proteomics reveals a global ubiquitination transition between 6 hr and 24 hr. (**A**) Experimental design for total protein and diGly peptides proteomic analysis by TMT labeling. (**B, C**) Volcano plots for the total protein abundance at 6 hr or 24 hr. Dashed lines represent the log_2_FC = ±1 and p-value = 0.01. The number of increased or decreased proteins (Log_2_FC > 1 or < -1, adj.p-value < 0.01) is indicated at right bottom corner. (**D, E**) Volcano plots for the diGly peptides abundance at 6 hr or 24 hr. Dashed lines represent the log_2_FC = ±1 and p-value = 0.01. The number of increased or decreased diGly peptides (Log_2_FC > 1 or < -1, adj.p-value < 0.01) is indicated at right bottom corner. (**F**) Venn diagram of proteins with increased ubiquitination in the diGly-TMT ubiquitinome and TUBE-TMT ubiquitinome at 6 hr post Poly (I:C) treatment. (**G**) Volcano plots for the diGly peptides abundance difference between 6 hr and 24 hr. Dashed lines and varying diGly peptide number are presented under same threshold as (D) and (E). (**H**) Venn diagram of the proteins having increased (or decreased) diGly peptides (Log_2_FC > 1 or < -1, adj.p-value < 0.01) in (G). (**I**) DiGly peptides abundance at different timepoints following Poly (I:C) treatment in THP-1 cells. (**J**) Ub linkage specific peptides portion in total diGly peptides at different timepoints. Statistical analysis was performed with one-way ANOVA method. *, p < 0.05; **, p < 0.01; ***, p < 0.001; ****, p < 0.0001; ns, not significant. (**K**) GO_ BP analysis of proteins with increased ubiquitination at Poly (I:C) 6 hr diGly-TMT proteomics. The top 20 biological processes were sorted by FDR and organized by Fold Enrichment. Red color represents the processes that are common to both timepoints. (**L**) GO_BP comparison between 6 hr and 24 hr. The top 20 processes were sorted by FDR or Fold Enrichment (FE). Details are presented in (K) and Figure S4. (**M**) List of the top 5 affected biological processes (selected by FE > 10, FDR < 0.01) at 6 and 24 hr post Poly (I:C) treatment.

We then focused on the diGly peptides abundance, which was normalized to the total protein abundance. We identified 12,285 diGly peptides, corresponding to 4,236 ubiquitinated proteins (table S4). The changes in diGly peptides abundance at the two timepoints were visually depicted in the volcano plots (Fig. 3D and E) and showed a significant increase at both 6 hr and 24 hr. These increased diGly peptides correspond to 1,009 and 1,032 proteins at 6 hr and 24 hr, respectively. These diGly-TMT protein sets represent the proteins with increased ubiquitination at 6 hr and 24 hr post Poly (I:C) treatment. Interestingly, the number of proteins with increased ubiquitination was much higher than those with decreased ubiquitination, and this observation explains the increase of global ubiquitination signal seen in immunoblotting (Fig. 1A). The comparison of these two sets showed that about one third of the proteins were unique to each timepoint (fig. S4A). Next, we compared this 6 hr proteins set to the previous TUBE-TMT identified 6 hr proteins with increased ubiquitination (Fig. 2C): 129 proteins, accounting for 59.5% of the TUBE-TMT proteomic identified, were shared by both approaches, revealing a substantial overlap (Fig. 3F).

To illustrate the ubiquitination transition between 6 hr and 24 hr, we compared the changes in diGly peptides abundance between these two timepoints. Unlike the dramatic increase shown in Figure 3D and E, we observed many diGly peptides’ abundance to be decreased at 24 hr when compared to 6 hr (Fig. 3G). The diGly peptides with increased or decreased abundance originated from 196 and 64 proteins, respectively (Fig. 3H). These two protein groups were different from each other, except for PRPF8, which had sites ubiquitinated and deubiquitinated simultaneously from 6 to 24 hr (Fig. 3H and fig. S4B). Next, we examined the variations in total diGly peptides amounts and Ub chains linkages. As shown (Fig. 3I), the amount of diGly peptides shifted significantly between 6 and 24 hr, reflecting a downregulation of the cellular ubiquitination state. In addition, the two main Ub chain linkages, K48 and K63, behaved differently at these two timepoints (Fig. 3J and fig. S4C). The K48 linkages decreased continuously from 0 to 24 hr; in contrast, the K63 linkages portion stayed nearly unchanged from 0 to 6 hr, and then increased slightly at 24 hr. Thus, the increased number of diGly peptides concomitant with a reduction in Ub chain linkages (fig. S4C) indicated that an extensive mono-ubiquitination of proteins took place.

The GO analysis of the proteins with increased ubiquitination revealed that changes in ubiquitination at 6 and 24 hr were associated with different biological processes (Fig. 3K and fig. S4D to F). Among the top 20 processes sorted by either false discovery rate (FDR) or Fold Enrichment (FE), 30% (6/20) or 40% (8/20) of the processes were exclusively identified at 6 hr or 24 hr post Poly (I:C) treatment respectively (Fig. 3L). For example, proteins involved in rRNA processing, cell cycle and apoptosis pathways were ubiquitinated at 6 hr, while proteins related to protein metabolism, lipid transport and MDA-5 signaling pathways were ubiquitinated at 24 hr (Fig. 3K and fig. S4D to F). To determine which processes were the most drastically ubiquitinated post Poly (I:C) treatment, we compared the top 5 processes (sorted by FE, with a threshold FE > 10 and FDR < 0.01) at both timepoints. Surprisingly, the top 5 processes were identical, and all related to intracellular protein localization, for example, protein localization to nucleoplasm (Fig. 3M). This indicates that Poly (I:C) induced increased ubiquitination regulates protein localization in cells.

In conclusion, our diGly-TMT proteomics analysis demonstrates that Poly (I:C) induces a dramatic increase in ubiquitination which does not lead to extensive protein degradation. Furthermore, a transition of Ub conjugates, including (1) the categories of ubiquitinated proteins, (2) the diGly peptides amount and (3) the types of Ub chain’s linkages, occurs between 6 and 24 hr, and this induced ubiquitination is involved in regulating intracellular protein localization.

### Influenza A virus infection upregulates global ubiquitination in monocytes

While Poly (I:C) is commonly used as a chemical mimic of viral RNA genomes, its mode of action differs slightly from actual viruses. Hence, we employed IAV as an RNA respiratory virus example to infect THP-1 cells and studied the IAV induced ubiquitinome. Initially, immunoblotting confirmed that IAV infection induced a similar upregulation of ubiquitination in THP-1 cells, although at a later timepoint (24 hr) as compared to Poly (I:C) treatment (Fig. 4A). This disparity may be attributed to the absence of the IAV entry and uncoating steps in Poly (I:C) transfection, leading to a more rapid effect. Consequently, we collected THP-1 cells at 0, 24, and 48 hr post-infection for ubiquitinome analysis with diGly-TMT mass spectrometry (table S5), accompanied with total protein abundance quantification and RNA sequencing (Fig. 4B).

**Fig. 4.**
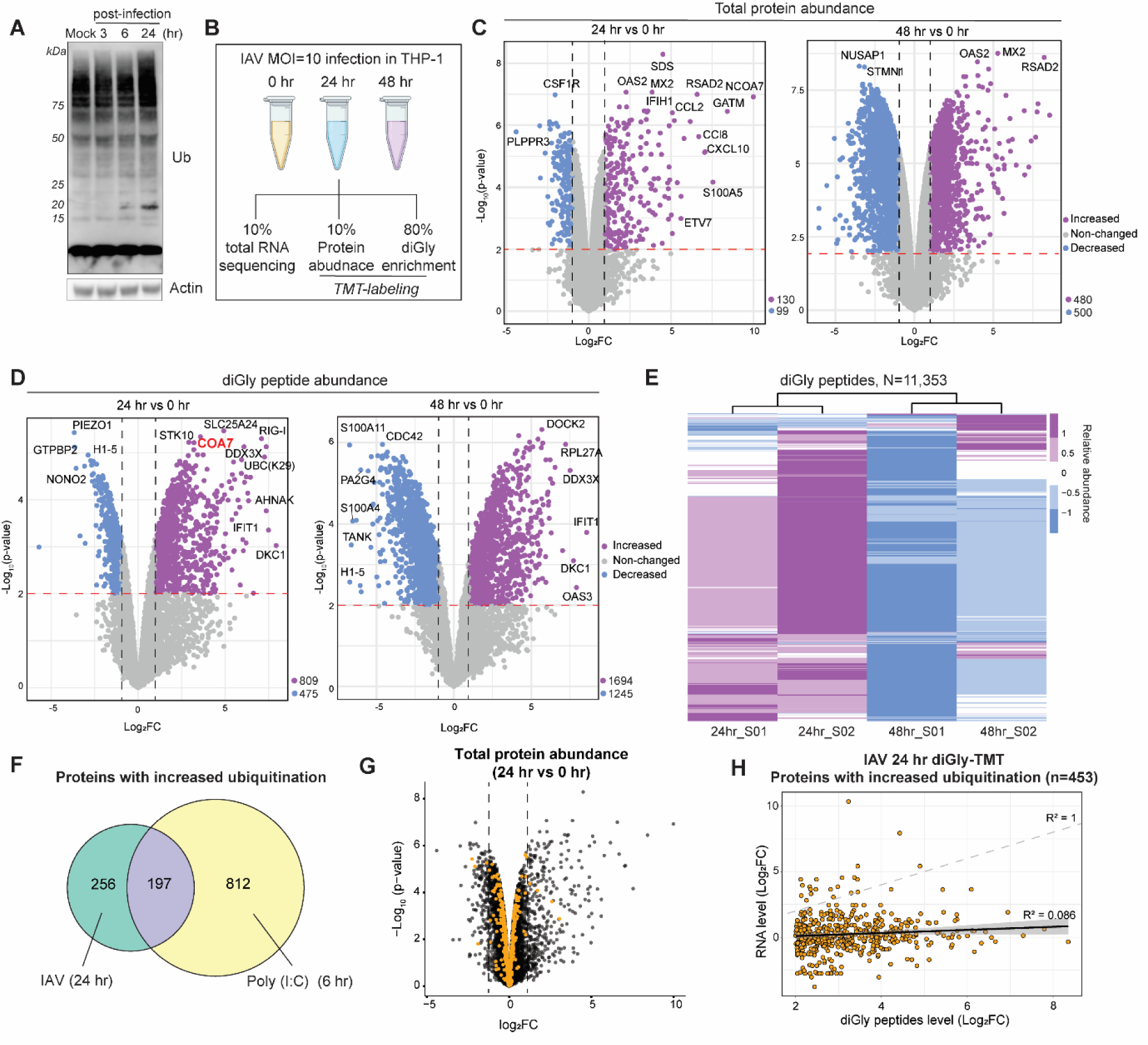
IAV induces a global ubiquitination increase at 24 hr in infected THP-1 cells. (**A**) Immunoblot for the global ubiquitination signal in IAV (X31, MOI = 10) infected THP-1 cells at 3, 6, and 24 hr post-infection. Actin was used as internal control. (**B**) Experimental design for the RNA sequencing and proteomic analysis of IAV infected THP-1 cells. (**C**) Volcano plots for the total protein abundance at 24 hr or 48 hr. Dashed lines represent the log_2_FC = ±1 and p-value = 0.01. The numbers of increased or decreased proteins (Log_2_FC > 1 or < -1, adj.p-value < 0.01) is indicated at right bottom corner. (**D**) Volcano plots for the diGly peptides abundance at 24 hr or 48 hr vs 0 hr. Dashed lines represent the log_2_FC = ±1 and p-value= 0.01. The number of increased or decreased diGly peptides (Log_2_FC > 1 or < -1, adj.p-value < 0.01) is indicated at right bottom corner. (**E**) Heatmap of diGly peptides abundance of the samples (S) in IAV infected THP-1 cells at 24 hr and 48 hr. S01 and S02 represent two biological replicates. (**F**) Venn diagram of proteins with increased ubiquitination following Poly (I:C) treatment (6 hr) or IAV infection (24 hr) in THP-1 cells. (**G**) Visualization of proteins with increased ubiquitination (n = 453) (orange dots) in total protein abundance volcano plot at 24 hr post-infection. Dashed lines represent the log_2_FC = ±1. (**H**) Alignment of diGly peptides to RNA level of the proteins with increased ubiquitination (n = 453) at 24 hr post IAV infection. Peptide and RNA level are presented as Log_2_FC (as compared to 0 hr timepoint).

Unlike Poly (I:C), virus infection led to a significant change in protein levels, especially at 48 hr post-infection (Fig. 4C). The variation in diGly peptides (Fig. 4D) was visualized after normalization to the protein abundance. Consistent with the immunoblotting findings in Figure 4A, at 24 hr post-infection diGly peptides with increased abundance (Log_2_FC > 1) (n = 809) were more than those with decreased abundance (Log_2_FC < -1) (n = 475), resulting in a notable upregulation of global ubiquitination. Later, at 48 hr post-infection, there were 1694 diGly peptides increased in abundance. Different from Poly (I:C), IAV infection led to a greater reduction in diGly peptides (n = 1245) at 48 hr (Fig. 4D). Then we compared the diGly peptides abundance of the 24 hr and 48 hr samples (Fig. 4E). As shown, there was a robust transition in ubiquitination as the abundance of most identified diGly peptides was generally reduced from 24 hr to 48 hr. This reduction may be connected to an induction of DUBs expression, as we found that the expression of some Ub specific peptidases (USPs), e.g., USP7, USP8 and USP18, was upregulated at 24 hr post-IAV infection (fig. S5A).

Since the onset of global ubiquitination increase was at 24 hr post-virus infection, we focused on the proteins with increased ubiquitination, which here we stringently defined as the proteins having at least one ubiquitinated peptide with a dramatic upregulation (Log_2_FC > 2 and adj.p-value < 0.05, details for calculating adj.p-value are in Methods “Statistical analysis” section). This resulted in the identification of 453 proteins (excluding isoforms, see table S6). 43% (197/453) of the proteins with increased ubiquitination at 24 hr had also been observed in the diGly-TMT analysis of 6 hr-Poly (I:C) treated cells (Fig. 4F). Considering that virus infection induced the expression of many genes and proteins (fig. S5B to D), we tested whether these increased ubiquitinated proteins were biased by changes in protein expression. First, after visualizing these 453 proteins in the “24 hr volcano plot” from Figure 4C, we found that the level of most proteins did not change significantly (Fig. 4G, -1 < Log_2_FC < 1). We also aligned the diGly peptides abundance variation to the RNA sequencing data and found that diGly peptides upregulation correlated poorly with RNA upregulation (Fig. 4H). Taken together, these data indicate that the IAV-induced protein’s ubiquitination upregulation is not biased by IAV-induced protein’s level changes.

Next, we quantified how many ubiquitinated peptides were found within each protein. By separating the total proteins into “proteins with increased ubiquitination at 24 hr (IAV)” and “other proteins”, we found that proteins with increased ubiquitination have a higher frequency of increased diGly peptides number (Fig. 5A). We also investigated whether multiple diGly peptides from the same protein were likely to be induced. To answer this, we considered all pairs of the 797 diGly peptides (from proteins with increased ubiquitination at 24 hr (IAV)) and recorded for how many of these pairs the two peptides came from the same protein (magenta line in Fig. 5B). We then repeated this for 1,000 random sets of 797 diGly peptides for comparison (grey histogram in Fig. 5B). As is visible, the fraction of pairs coming from the same protein was considerably larger among proteins with increased ubiquitination. These data indicate that once a protein is ubiquitinated under IAV infection, it tends to have more than one ubiquitinated site. Next, we examined the diGly peptides abundance variation (Fig. 5C). As expected, there was a significant increase in the number of diGly peptides at 24 hr compared to 0 hr, and at 48 hr, the number of diGly peptides was reduced (compared to 24 hr). Finally, we investigated how the Ub chain linkages varied during IAV infection. Like Poly (I:C) treatment, the abundance of K48 linkages decreased several fold from 0 to 48 hr, while K63 linkages stayed unchanged (Fig. 5D). Surprisingly, another minor linkage, K29, increased dramatically at 48 hr as compared to both 0 and 24 hr, unlike what had been observed in the Poly (I:C) treatment (fig. S4C). In summary, we found that the IAV-induced ubiquitination increase is specific. Moreover, between 24 and 48 hr, ubiquitination shows an abrupt transition, with large changes in diGly peptides abundance and chain linkages composition.

**Fig. 5.**
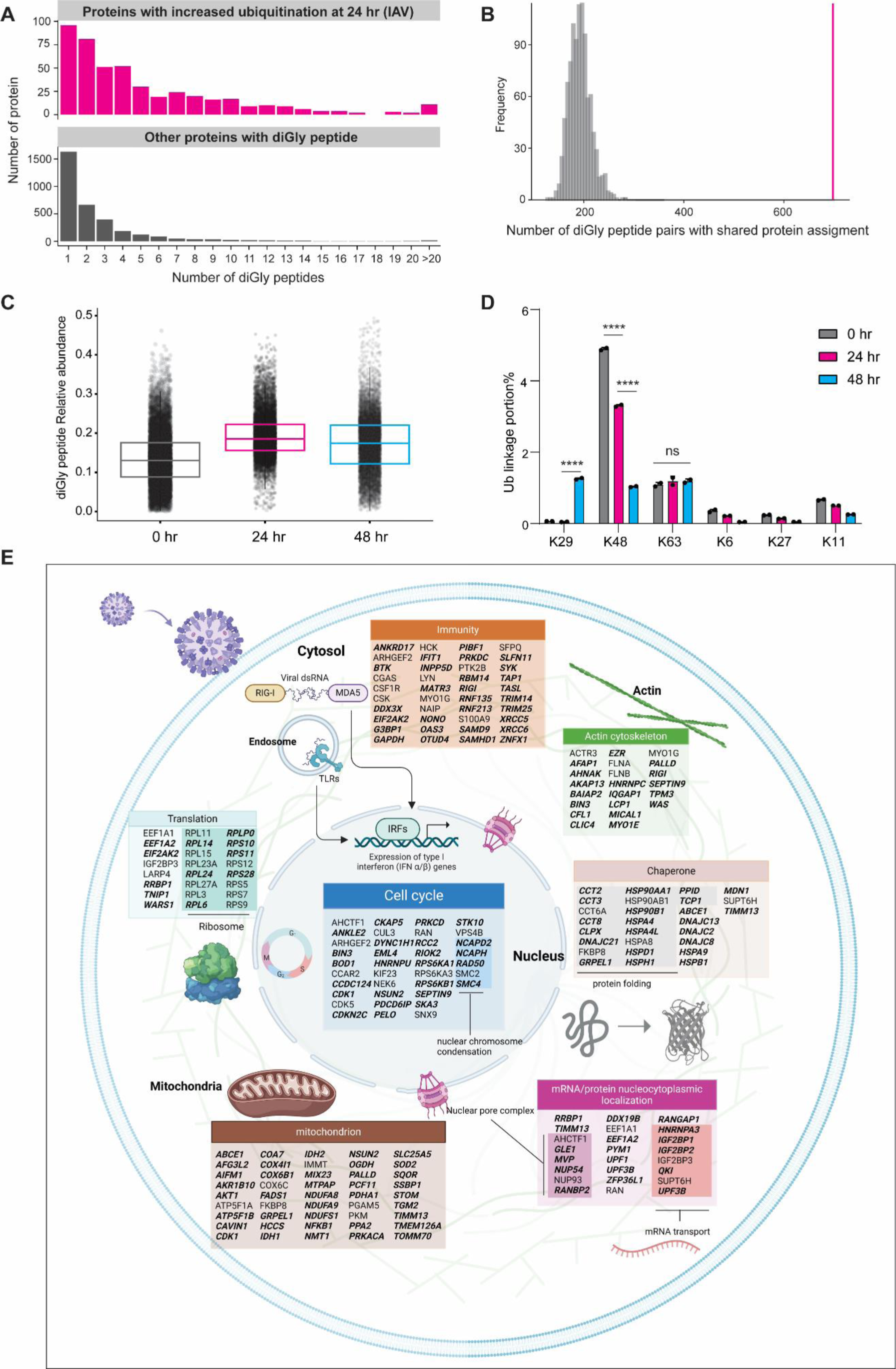
IAV induced ubiquitination in THP-1 cells. (**A**) Analysis of the number of ubiquitinated peptides per protein at 24 hr post IAV infection. (**B**) Analysis of randomly paired ubiquitinated peptides in shared protein. The magenta line represents the value from proteins with increased ubiquitination at 24 hr (IAV), and the grey histogram represents the value from randomly chosen 1,000 sets of 797 diGly peptides. (**C**) DiGly peptides abundance at all 3 timepoints under IAV infection in THP-1 cells. (**D**) Ub linkage specific peptides portion in total diGly peptides at all 3 timepoints under IAV infection in THP-1 cells. Statistical analysis was performed with one-way ANOVA test. *, p < 0.05; **, p < 0.01; ***, p < 0.001; ****, p < 0.0001; ns, not significant. (**E**) DAVID clustering analysis of all the IAV induced ubiquitinated proteins (with diGly peptides abundance Log_2_FC > 1, adj.p-value < 0.01) at 24 hr. Each cluster was chosen by FDR (< 0.01). Proteins with increased ubiquitination (n = 453, in Fig. 4F) are bold and italic.

To better understand the role of the ubiquitinated proteins at 24 hr post IAV infection, we first performed the GO analysis for the “proteins with increased ubiquitination at 24 hr (IAV)”. We found that protein localization processes were strongly enriched (fig. S5E), consistent with our previous findings with Poly (I:C) treatment (Fig. 3M). Later, we analyzed the ubiquitinated proteins (diGly peptide Log_2_FC > 1) for their biological function using DAVID clustering. The proteins were allocated into several major clusters representing different biological processes, including mitochondria and protein/mRNA localization (Fig. 5E). Notably, many proteins connected to protein localization in the nucleus were found to be ubiquitinated under IAV infection. Among them, components of the nuclear pore complex, including MVP, RanBP2, NUP54, NUP93 and AHCTF1, were identified as a major protein complex regulated by ubiquitination. This observation indicates that ubiquitination may modulate protein localization during IAV infection.

Next, to investigate the IAV induced ubiquitination in a more physiological sample, we infected mice with IAV intra-tracheally (*48*) and 7 days later isolated *in vivo* monocytes by multi-color staining and FACS sorting (fig. S6A and B). Due to the limited number of *in vivo* monocytes isolated, conventional ubiquitinome analysis was not possible. We therefore used a protocol similar to that used in single-cell proteomics and employed a carrier TMT channel to characterize ubiquitinated proteins (*49*). In detail, for each individual TMT channel, 0.3 million FACS sorted cells (obtained from 2 mice) were used. For the carrier channel, 11 million cells from both non-infected and infected mice were utilized. Total protein abundance was analyzed additionally (fig. S6C). First, as depicted in fig S7A, infected mice experienced continuous weight loss until day 7, indicating successful infection. Flow cytometry analysis of total lung cells showed an enrichment of proinflammatory CD45+CD11b+Ly6G-Ly6C^High(Hi)^ monocytes as reported (*35, 48, 50*). The proportion of Ly6C^Hi^ monocytes increased from 1% to 10%, while Ly6C^Low(Lo)^ monocytes remained almost unchanged (fig. S7B and C). The IAV induced enrichment of Ly6C^Hi^ monocytes motivated us to explore their ubiquitinome. In comparison to our previous proteomic analyses, only few diGly peptides were identified (fig. S7D), and the expression level of many proteins changed significantly in response to IAV infection (fig. S7E). After normalizing diGly peptides abundance to protein abundance, we merely obtained 8 diGly peptides with significant increase in abundance (fig. S7F and table S7); most of them had been identified in the proteins with increased ubiquitination in 6 hr Poly (I:C) treated or 24 hr IAV infected THP-1 cells (fig. S7G).

### Ubiquitination leads to degradation of EIF4H under IAV infection

Next, to explore the function of viral RNA or IAV induced ubiquitination, we focused on the ubiquitinated proteins identified in both Poly (I:C) treatment and IAV infection. Considering that our proteomic experiments had been performed in cells with an active UPS (i.e. without inhibition of the proteasome), it was possible to investigate proteins that were ubiquitinated and degraded in response to Poly (I:C) treatment and virus infection. Such proteins should show a significant reduction in protein abundance and be positive for diGly peptides. Although the abundance of most proteins remained unchanged following Poly (I:C) treatment, a subset was degraded. Since translation initiation factors have been shown to be ubiquitinated under heat-shock (*20*), and viral RNA can manipulate cellular translation (reviewed in (*51*)), we first looked at the abundance of eukaryotic initiation factors (EIFs). Total protein abundance analysis at both 6 and 24 hr revealed a markedly lower EIF4H level (Fig. 6A and table S3) in Poly (I:C) treated THP-1 cells. Immunoblotting confirmed this reduction in both THP-1 (Fig. 6B) and A549 (Fig. 6C and fig. S8) cells. Notably, addition of the proteasome inhibitor Bortezomib restored EIF4H levels, indicating that the degradation is proteasome and ubiquitination dependent.

**Fig. 6.**
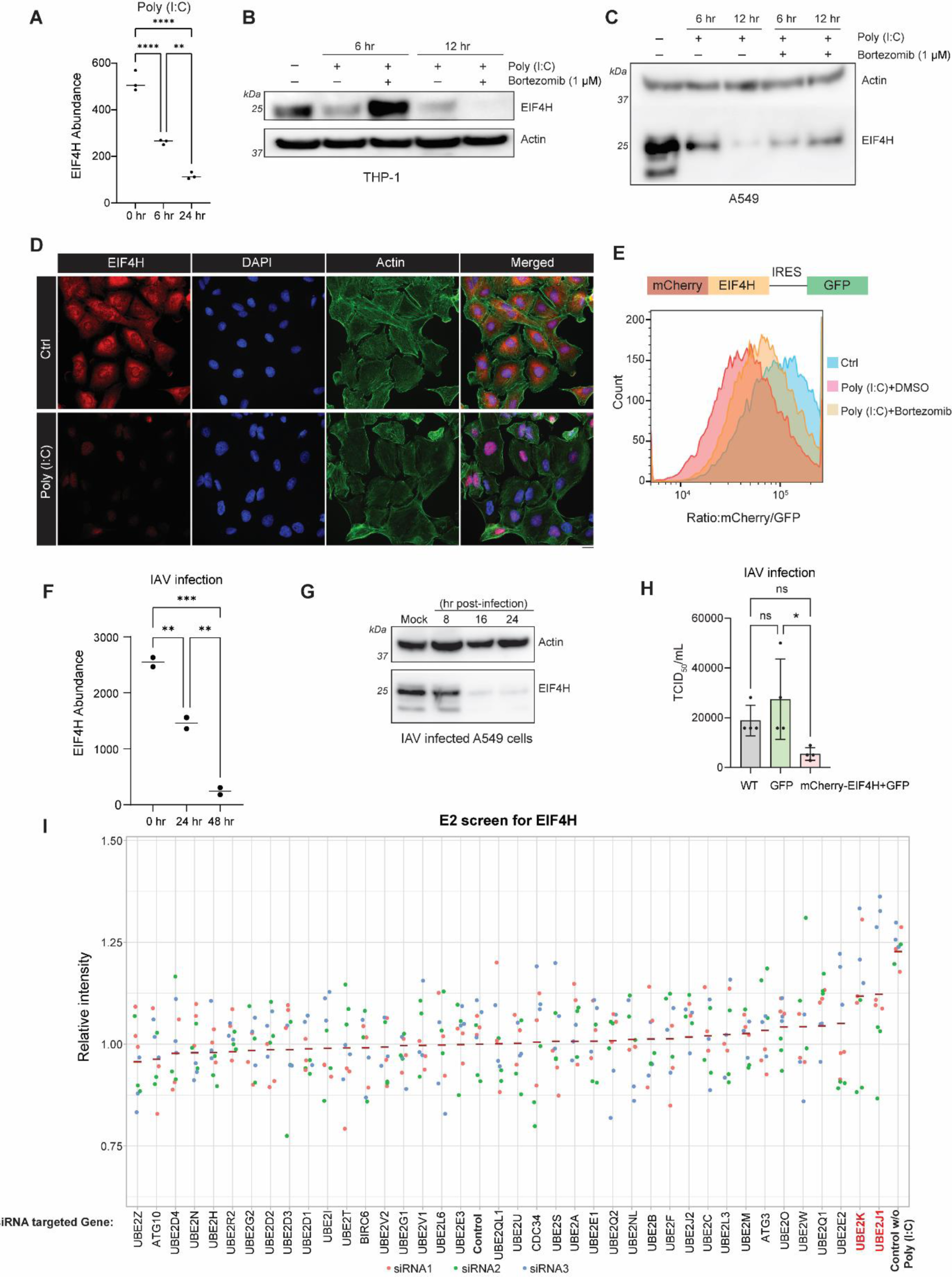
Proteolytic ubiquitination of EIF4H. (**A**) EIF4H abundance in total protein analysis in Poly (I:C) treated THP-1 cells. Statistical analysis was performed with one-way ANOVA test. **, p < 0.01; ****, p < 0.0001. (**B**) Immunoblotting of EIF4H in Poly (I:C) (2.5 μg/mL) treated THP-1 cells. Bortezomib (1 μM) was added together with Poly (I:C) treatment, time as indicated. Actin was used as internal control. (**C**) Immunoblotting of EIF4H in Poly (I:C) treated A549 cells. Poly (I:C) concentration was 5 µg/mL, other inhibitors as in (B), time as indicated. (**D**) Immunofluorescence of EIF4H in Poly (I:C) treated A549 cells. Poly (I:C) concentration was 5 µg/mL for 12 hr. Scale bar is 10 μm. (**E**) FACS analysis of mCherry-EIF4H degradation in A549 cells stably expressing mCherry-EIF4H-IRES-GFP. Poly (I:C) treatment was 5 μg/mL for 24 hr. Bortezomib (1 μM) was added together with Poly (I:C). (**F**) EIF4H levels in total protein abundance analysis in IAV infected THP-1 cells. Statistical analysis was performed with one-way ANOVA test. **, p < 0.01; ***, p < 0.001. (**G**) Immunoblotting of EIF4H in IAV (X31, MOI = 1) infected A549 cells from 8 hr to 24 hr post-infection. (**H**) TCID_50_ assay to measure IAV titers from the supernatant of IAV-infected cell lines (n = 4). IAV X31 with MOI = 0.1 was used for infection. Supernatant was collected at 48 hr post-infection. Statistical analysis was performed with one-way ANOVA test. *, p < 0.05; ns, not significant. (**I**) E2 ligase screen for EIF4H. A549 cells were first reverse transfected with different E2 siRNAs and then treated with Poly (I:C) (5 μg/mL, 12 hr) to induce EIF4H degradation. EIF4H level was measured by immunofluorescence and normalized to control group. 3 different siRNAs were designed for each E2 ligase. Control: A549 cells were transfected with non-targeting siRNAs and then treated with Poly (I:C); Control w/o Poly (I:C): A549 cells were transfected with non-targeting siRNAs only.

We further investigated EIF4H degradation using immunofluorescence and flow cytometry. In fixed control A549 cells staining with an antibody against EIF4H showed a homogeneous pattern across the cytoplasm and the nucleus. In Poly (I:C) treated cells, the staining was abolished, indicating that most of the endogenous EIF4H was degraded (Fig. 6D). Next, we established a reporter cell line overexpressing EIF4H as a fusion protein to mCherry: the construct also contained an internal ribosome entry site (IRES) and GFP as an internal control (Fig. 6E, upper). Expression of EIF4H-mCherry and GFP signals was monitored by flow cytometry. As shown in Figure 6E, Poly (I:C) led to a strong decrease in mCherry/GFP ratio, indicating EIF4H degradation. Co-administration of Bortezomib largely prevented the mCherry signal loss. Finally, we verified the impact of IAV infection on EIF4H protein levels in THP-1 cells. As visible in Figure 6F, EIF4H levels dropped dramatically at 24 hr post-infection and continued to decrease at 48 hr. This reduction was also observed in IAV-infected A549 cells (Fig. 6G), where EIF4H protein levels significantly diminished from 16 hr post-infection. Given that EIF4H degradation was induced by IAV, we hypothesized that EIF4H may be an antiviral host protein. To test this, the viral titer was measured in IAV-infected cells. As shown in Figure 6H, EIF4H overexpressing cells had a lower viral titer in the supernatant compared to both wild-type (WT) and GFP overexpressing A549 cells.

Finally, we used an image-based screen assay to identify the E2 ligase responsible for EIF4H ubiquitination. Various E2 ligases were first knocked down by siRNA, and then cells were treated with Poly (I:C) for 12 hr. After that, EIF4H intensity was visualized by immunofluorescence and normalized to the control group (Fig. 6I). As shown, knockdown of UBE2K and UBE2J1 restored EIF4H intensity in Poly (I:C) treated A549 cells, suggesting an inhibition on EIF4H’s ubiquitination. Interestingly, both E2s can mediate endoplasmic reticulum-associated degradation (ERAD) of abnormal proteins (*52, 53*), and especially for UBE2J1, it is essential for cells to recover from ER stress (*53*). Overall, our data suggest that EIF4H, is a potentially antiviral protein, that is ubiquitinated by UBE2K and UBE2J1, and then degraded by the proteasome following IAV infection, thereby facilitating virus infection.

### Non-proteolytic ubiquitination of COA7 leads to its nuclear localization

Unlike EIF4H, for most proteins, Poly (I:C) induced ubiquitination rarely affected their abundance (Fig. 3B and C). This suggests a non-proteolytic role of ubiquitination in regulating the cellular stress response against viral RNA challenge. In this context, we studied the mitochondrial Fe-binding protein COA7 (*54*), one of the top ubiquitinated hits from our proteomic data in both Poly (I:C) treatment and IAV infection, with unchanged protein abundance. Its diGly peptides revealed several sites ubiquitinated under either Poly (I:C) treatment or IAV infection (Fig. 7A). Notably, K56 and K90, which were ubiquitinated under both treatments, are located on the surface of COA7 suggesting that they might be involved in regulating interactions with other proteins (fig. S9A and B). Due to the sensitivity of THP-1 cells to exogenous RNA treatment and their limited suitability for immunofluorescence studies, we used A549 cells for the following experiments. We first confirmed COA7’s ubiquitination in A549 cells. After TUBE pulldown, a COA7 signal was detected under Poly (I:C) treatment, which disappeared with Bafilomycin A1 or TAK-243 addition (Fig. 7B).

**Fig. 7.**
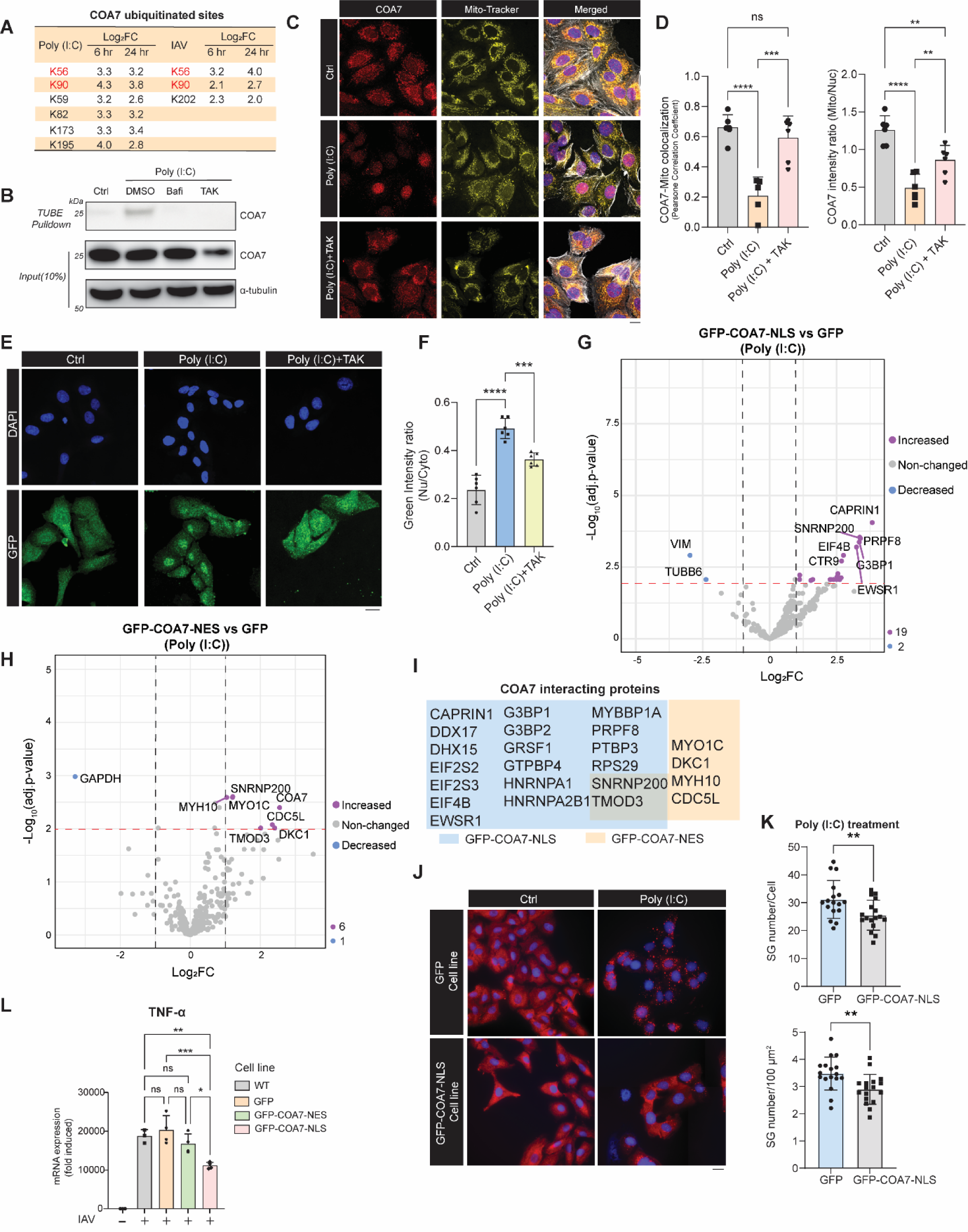
Non-proteolytic ubiquitination of COA7. (**A**) Induced ubiquitinated sites (Log_2_FC > 1, adj.p-value < 0.01) in COA7 under Poly (I:C) treatment or IAV infection. Sites which are shared by both treatments are colored in red. (**B**) Verification of COA7 ubiquitination by TUBE pulldown in Poly (I:C) treated A549 cells. Poly (I:C) treatment was 5 ug/mL for 6 hr. Bafilomycin A1 (500 nM) and TAK-243 (1 μM) were added together with Poly (I:C) transfection. (**C**) Immunostaining of COA7 in A549 cells under Poly (I:C) and TAK-243 treatment. Poly (I:C) treatment was 5 ug/mL for 6 hr, and TAK-243 (1 μM) was added together with Poly (I:C) transfection. Scale bar is 20 μm. (**D**) Quantification of COA7/Mito-Tracker colocalization and the ratio of mitochondrial COA7 to nuclear COA7. Statistical analysis was performed with one-way ANOVA test. *, p < 0.05; **, p < 0.01; ***, p < 0.001; ****, p < 0.0001; ns, not significant. 6 pictures from the 40x objective were used, and in total > 200 cells for each group were quantified. (**E**) GFP signal of GFP-COA7-NES cell line under Poly (I:C) and TAK-243 treatment (as in (C) above). Scale bar is 20 μm. (**F**) Ratio of nuclear green intensity to cytosolic green intensity in (E). Statistical analysis was performed with one-way ANOVA test. ***, p < 0.001; ****, p < 0.0001. 6 pictures from 40x objective were used, and in total > 200 cells for each group were quantified. Nu: nuclear; Cyto: cytosolic. (**G, H**) Volcano plots for NLS-COA7 and NES-COA7 interacting partners under Poly (I:C) treatment. Dashed lines represent the log_2_FC = ±1, and adj.p-value = 0.01. The number of increased and decreased proteins (Log_2_FC > 1 or < -1, adj.p-value < 0.01) is indicated at right bottom corner (COA7 was excluded). (**I**) Summary of COA7 interacting proteins (Log_2_FC > 1, adj.p-value < 0.01) from (G) and (H). (**J**) Stress granules formation in COA7-NLS cell line under Poly (I:C) treatment (5 μg/mL for 6 hr). Red signal represents the G3BP1 intensity. Blue (DAPI) signal represents the nucleus. Scale bar is 20 μm. (**K**) Quantification of stress granules formation in (J). Statistical analysis was performed with unpaired T-test. p< 0.05 was considered as significant. **, p < 0.01. >10 pictures from 40x objective were used, and in total > 400 cells for each group were quantified. (**L**) RT-qPCR of 24 hr IAV infection (X31, MOI = 1) induced TNF-α expression in different A549 cell lines (n = 3). The 2^-ΔΔct^ method was used for relative quantification. GAPDH was used as reference gene. Statistical analysis was performed with one-way ANOVA test. *, p < 0.05; **, p < 0.01; ***, p < 0.001; ns, not significant.

Given the limited information on COA7, we first employed siRNA for knock-down experiments (fig. S9C) and monitored the expressions of inflammatory and antiviral genes. This resulted in a significant upregulation of TNF-α, while expression of IFNB1, IFIT2 and type III interferon IFNL1 was little affected (fig. S9D). We then overexpressed COA7 by lentivirus transduction and observed a reduction in TNF-α upregulation following Poly (I:C) treatment (fig. S9E), indicating that COA7 may act as a repressor for TNF-α expression.

COA7 is a cytochrome c assembly factor, and cytochrome c is known to translocate from mitochondria to the nucleus during apoptosis (*55, 56*). Our BP analysis presented above indicates that both Poly (I:C) and IAV induced ubiquitination are involved in protein nucleocytoplasmic localization (Fig. 3M and Fig. 5E). Therefore, we examined COA7’s localization by immunofluorescence. In the control condition, COA7 was found in mitochondria as expected, but after 6 hr Poly (I:C) treatment or 24 hr post IAV infection, its intensity in mitochondria significantly decreased (Fig. 7C and D and fig. S9F). Co-treatment with TAK-243 restored the mitochondrial COA7 signal, indicating a ubiquitination-dependent subcellular localization. To confirm the re-localization, we prepared COA7–GFP fusion proteins containing either a nuclear export signal (NES) or a nuclear localization signal (NLS) and established stable expressing cell lines (fig. S9G). In the GFP-COA7-NES cell line, the signal in the control condition was cytoplasmic, and broadly similar to the endogenous COA7. Poly (I:C) treatment induced a strong enrichment of the GFP signal in the nucleus (Fig. 7E) and an increase in the nuclear GFP/cytosolic GFP ratio (Fig. 7F), both of which were inhibited by TAK-243 addition.

To understand the consequence of COA7 translocation, we used the GFP-COA7-NES and GFP-COA7-NLS cell lines to determine the interactome of cytosolic (cy) and nuclear (nu) COA7, respectively (table S8). As shown in Figure 7G and H, COA7-NES (cyCOA7) and COA7-NLS (nuCOA7) interacted with different proteins (represented as purple dots with Log_2_FC > 1, adj.p-value < 0.01). nuCOA7, but not cyCOA7, was involved in SGs pathway as the core components of SGs, e.g., CAPRIN1, G3BP1 and G3BP2, were enriched in GFP-COA7-NLS cells but not in GFP-COA7-NES cells (Fig. 7I). Absence of Poly (I:C) or addition of TAK-243 strongly reduced the interaction (fig. S9H and I). Therefore, we examined SGs formation in the COA7-NLS cell line. As expected from the above results, nuCOA7 inhibited Poly (I:C)-induced SGs formation (Fig. 7J and K), while arsenite-induced SGs were not affected (fig. S9J and K). Since G3BP1 is present in both Poly (I:C)- and arsenite-induced SGs (*57-59*), we reasoned that nuCOA7, like ubiquitinated and translocated COA7, may impact the RNA transport process.

Additionally, we probed the impact of COA7 localization on IAV infection. The different cell lines were infected with IAV and after 24 hr RNA was extracted and examined for the expression of inflammatory and antiviral genes. The RT-qPCR result revealed that nuCOA7 partly inhibited TNF-α expression, while cyCOA7 did not (Fig. 7L). Moreover, IFNB1’s expression showed no difference between nuCOA7 and cyCOA7 (fig. S9L). In conclusion, under Poly (I:C) treatment or IAV infection, COA7 undergoes ubiquitination, localizes to the nucleus, and this impacts Poly (I:C) induced SGs formation and IAV-induced TNF-α expression.

### Ubiquitination regulates viral protein and viral RNA translocation

By focusing on the host cell protein COA7, we found that ubiquitination is critical for its cytosolic or nuclear localization. We then investigated whether IAV induced ubiquitination also regulates the localization of IAV proteins. To this end, we monitored the distribution of the viral nucleoprotein (NP) during IAV infection. As shown in Figure 8A and B, at 10 hr post-infection, NP was widely distributed in infected A549 cells, especially in the cytosol. However, when cells were treated together with TAK-243, NP was restricted to the nucleus. Given that viral RNA assembles with NP into viral ribonucleoproteins (vRNPs), we also examined the localization of double-stranded (ds) viral RNA. After blocking ubiquitination, like NP, ds viral RNA was also found mostly in the nucleus, whereas under normal conditions, it was distributed throughout the cell (Fig. 8C and D).

**Fig. 8.**
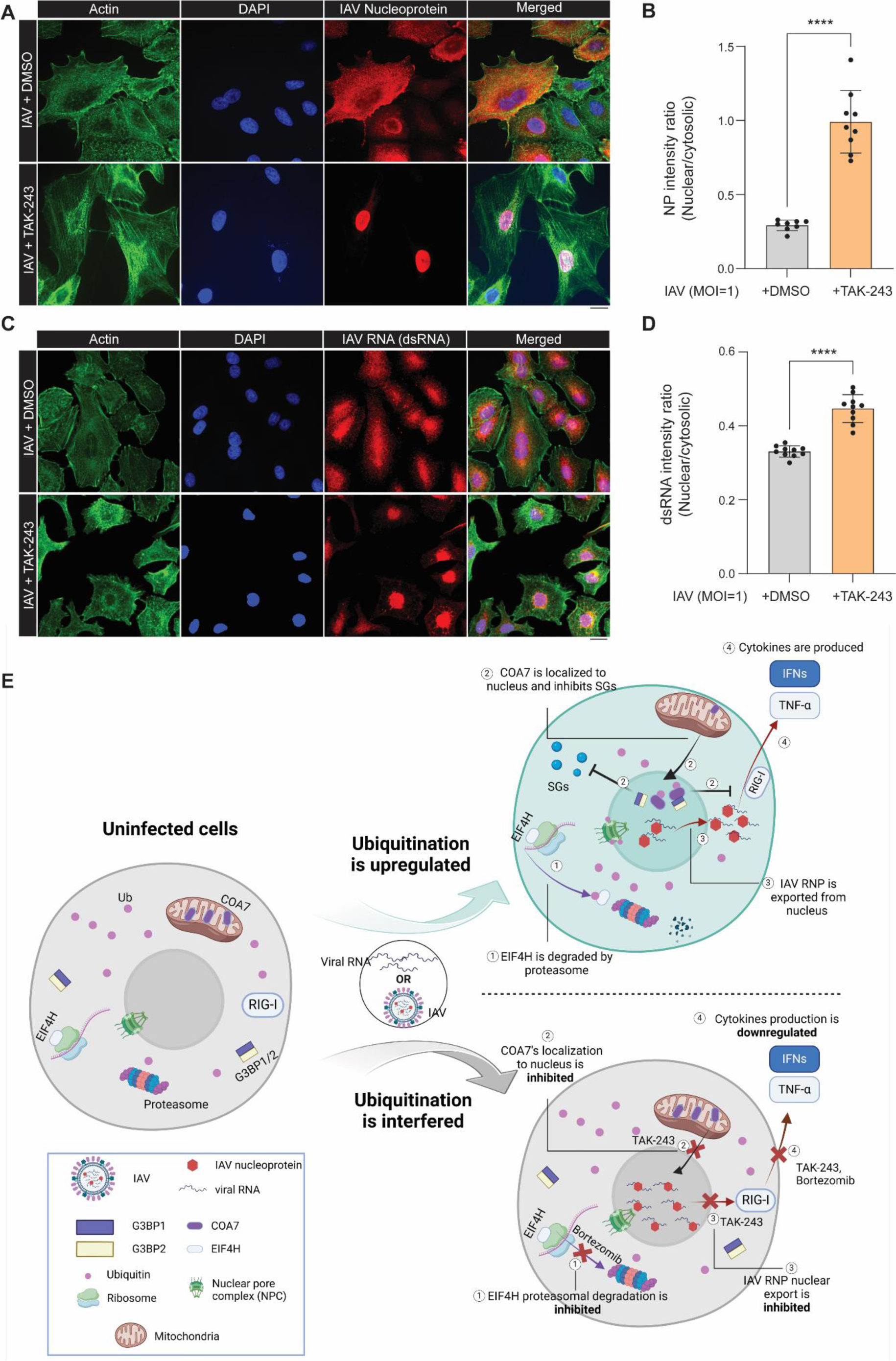
Ubiquitination regulates viral protein and dsRNA localization under IAV infection. (**A**) Immunostaining of IAV NP protein (red) in IAV (X31, MOI = 1) infected A549 cells at 10 hr post-infection. Scale bar is 20 μm. (**B**) Quantification of the ratio (nuclear NP intensity/cytosolic NP intensity). 9 pictures from 40x objective were used, and in total > 200 cells were quantified. Statistical analysis was performed with unpaired T-test. ****, p < 0.0001. (**C**) Immunostaining of IAV dsRNA (red) in IAV (X31, MOI = 1) infected A549 cells at 10 hr post-infection. Scale bar is 20 μm. (**D**) Quantification of the ratio (nuclear dsRNA intensity/cytosolic dsRNA intensity). 10 pictures from 40x objective were used, and in total > 200 cells were quantified. Statistical analysis was performed with unpaired T-test. ****, p < 0.0001. (**E**) Proposed model illustrating the functions of viral RNA or IAV infection induced global ubiquitination. Affected processes are listed as ①, ②, ③, and ④.

Based on the above findings, we propose a working model for viral RNA-induced global ubiquitination increase in monocytes (Fig. 8E). When cells are challenged with viral RNA mimics like Poly (I:C) or RNA virus infection by IAV, there is a notable rise in global ubiquitination (at 6 hr for Poly (I:C) treatment and 24 hr for IAV infection). This induced ubiquitination is widespread and not primarily proteolytic. We further propose that it may impact several processes related to virus infection and lead to: (1) Degradation of antiviral proteins. Many proteins undergo ubiquitination, and for some, like the antiviral protein EIF4H, ubiquitination leads to its degradation (Fig. 4D), facilitating IAV infection. (2) Regulation of subcellular localization. For other proteins, such as the mitochondrial protein COA7, ubiquitination regulates its subcellular localization, causing a translocation from the mitochondria to the nucleus. The ubiquitinated and translocated COA7 interacts with SGs core components G3BP1 and G3BP2 and inhibits viral RNA-induced SGs formation. Additionally, nuclear COA7 suppresses IAV-induced TNF-α expression. (3) IAV RNP’s nuclear export. Ubiquitination also determines the export of IAV NP and vRNA from the nucleus. Inhibiting ubiquitination with TAK-243 during IAV infection confines NP and viral RNA to the cell nucleus and inhibits the expression of TNF-α and IFN-β. (4) Proinflammatory and antiviral cytokines release. After viral RNA challenge, interfering with ubiquitination at different steps by various inhibitors strongly inhibits proinflammatory and antiviral cytokines release, e.g., TNF-α and IFN-β.

## Discussion

Our proteomic and biochemical results reveal a viral RNA and IAV induced global ubiquitination increase and explore both the proteolytic and non-proteolytic functions of this induced ubiquitination. Interestingly, an increase in ubiquitination was observed by immunoblotting in monocytes like THP-1 and BMDMs, but not in the epithelial cell line A549. This indicates that virus infection induced global ubiquitination increase may be a cell type-specific response. Yet, the lack of robust ubiquitination upregulation in IAV-infected A549 cells does not necessarily imply that ubiquitination is not modulated in these cells, as it was recently shown that SARS-CoV-2 can induce the ubiquitination of hundreds of proteins in A549 cells (*27*). Therefore, the modulation of ubiquitination under virus infection likely depends on the virus and cell type. Furthermore, it is possible that in our experiments with IAV infected A549 cells there were dynamic changes in ubiquitination, but the cells maintained a balance between ubiquitination and deubiquitination, so that no visible increase was observed in immunoblots. Considering the distinct roles of monocytes and epithelial cells during IAV infection, we propose that the increased ubiquitination in monocytes may be crucial for the release of various proinflammatory and antiviral cytokines. This hypothesis is supported by our results (Fig. 1C and fig. S1C to G), which show that interfering with different steps of the ubiquitination process significantly inhibits the expressions of TNF-α, and other antiviral factors like IFN-β, IFIT2, and RIG-I/DDX58.

In addition, we observed that the cellular localization of both host cells and viral proteins is determined by ubiquitination. Given that viral proteins produced by the host cell machinery must be transported and assembled into functional viral particles during infection, ubiquitination-mediated protein transport may be crucial for successful virus infection. There are two possible mechanisms by which ubiquitination can regulate protein transport: (1) Ubiquitination of transport complexes: under stress conditions, protein transport complexes (e.g., nuclear pore complex) or related proteins become ubiquitinated, affecting their functions and modulating localization (*60, 61*). This is supported by our findings that many proteins involved in transport such as MVP and RanBP2 are among the proteins with increased ubiquitination under IAV infection (Fig. 5E). (2) Ubiquitination of transported proteins: the cargo proteins themselves might be ubiquitinated, altering their interactions with transport complexes and further impacting their subcellular localization (*62, 63*). We cannot exclude either of these mechanisms, and both may play a role in stress-induced ubiquitination. Previous studies have shown that heat-shock-induced ubiquitination regulates protein transport between the nucleus and the cytosol (*20, 64*); while the mechanisms involved have not been defined, it was shown in another study that SGs assembly disrupts nucleocytoplasmic transport (*65*).

Although the global ubiquitination level increased after Poly (I:C) treatment or IAV infection, the various types of Ub linkages representing different Ub chains behave differently. As shown in Figure 3J and 5D, we observed a significant reduction in K48 linkage and no change in K63 linkage, indicating a reduction of the average length of Ub chains. While a significant increase in K29 linkage was observed at 48 hr post-IAV infection, its overall contribution was too low to compensate for the reduction in K48 linkage. Moreover, K29 linkage was not increased, neither at 6 hr after Poly (I:C) treatment nor at 24 hr post IAV infection. Given the increased total diGly peptides amount after Poly (I:C) treatment at 6 hr (Fig. 3I) and IAV infection at 24 hr (Fig. 5C), we surmise that the upregulation of global ubiquitination at early stage is mostly due to the increased mono-ubiquitination, which leads to a higher abundance of diGly peptides but not to an increase in linkage-specific peptides. This hypothesis also partially explains why fewer proteins are identified in TUBE pulldown compared to diGly enrichment, as TUBE likely has higher baseline affinity for binding poly-ubiquitinated over mono-ubiquitinated proteins (*20, 66*). Thus, to gain a detailed understanding of the cellular ubiquitinome, it is important to use a combination of approaches.

Mono-ubiquitination is strongly associated with the non-proteolytic functions of ubiquitination (*67*). For example, in response to DNA damage, PCNA undergoes mono-ubiquitination (*68, 69*) and interacts with Y family DNA polymerases (*70, 71*). Mono-ubiquitination also regulates protein localization, as initially observed in yeast, where it determines plasma protein internalization (*72, 73*). Additionally, the transcription factor p53 is exported out of the nucleus following its mono-ubiquitination, which increases the accessibility of its nuclear export sequence (*74*). Unlike the irreversible proteolytic ubiquitination of p53, mono-ubiquitination can be reversed by the deubiquitinase USP10, allowing the reimport of p53 into the nucleus (*75*). We hypothesize that the increased mono-ubiquitination in monocytes might be a temporary state induced by RNA virus infection. This increase in global mono-ubiquitination may affect protein solubility, interactions, localization, and enzymatic activity to cope with the infection, allowing the cell to adapt to pathogenic stress. Once the infection is stopped, mono-ubiquitination levels may revert to normal.

Finally, there is a seeming paradox between the global ubiquitination increase and the presumed specificity of Ub ligases. Traditionally, E3 ligases transfer Ub from E2 enzymes to target proteins by recognizing specific degrons or interacting with distinct protein domains (reviewed in (*76*)). These mechanisms rely on 3D structure recognition, implying high selectivity for E3 ligases. However, it remains possible that one or a few E3 ligases with low substrate selectively may be responsible for the global ubiquitination increase, a phenomenon that has been reported for ISGlyation by HERC6 (*77*). Additionally, considering that E2 enzymes can interact with multiple E3 ligases, it is plausible that one or several E2 enzymes are the primary mediators of the viral RNA-induced global ubiquitination increase. Identifying these key E2 or E3 ligases could pave the way for the development of clinically valuable inhibitors, presenting new strategies for antiviral and anti-inflammatory therapies.

## Acknowledgements

We thank Laure Plantard and Laurent Gelman for setting up and helping with the microscopy experiments. We thank Daniel Hess and Jan Seebacher for helping with the mass spectrometry experiment. We thank FMI functional genomic facility for sequencing the IAV infected samples. We thank Hans-Rudolf Hotz and Alan Naylor for helping upload the genomic and proteomic data to publica database. The working model was created with BioRender.com.

## Authors contributions

Conceptualization: LW, PM

Methodology: SS, LW, VI, ASS, HK, JC, SG, CC

Investigation: SS, LW, VI, CS

Visualization: SS, LW, VI, ASS, CS

Funding acquisition: YY, PM

Project administration: GM, PM

Supervision: YH, YY, PM

Writing – original draft: SS, PM, LW

Writing –editing: SS, VI, CS, YY, PM, LW

## Data and materials availability

All unique reagents generated in this study are available from the lead contact with a completed Materials Transfer Agreement (MTA); published research reagents from the FMI are shared with the academic community under an MTA having terms and conditions corresponding to those of the UBMTA (Uniform Biological Material Transfer Agreement).

RNA sequencing data have been deposited at ArrayExpress (accession E-MTAB-14283) and are publicly available as of the date of publication. Mass spectrometry data (Search results are provided as supplemental tables) have been deposited at the ProteomeXchange Consortium via the PRIDE partner repository and are publicly available as of the date of publication. For access the original proteomic data, information is in following: PXD054038 (Token: rsuAiuFqtvBj) is related to Figure 1; PXD054047 (Token: GQoEnrv56ZfL) is related to Figure 2; PXD054104 (Token: SLbUXPXJanQw) and PXD054106 (Token: I0GGSkzsbTBf) are related to Figure 3; PXD054110 (Token: 53vmUE7vguLo) and PXD054111 (Token: fi1AJg2Q9Vye) are related to figure S7; PXD054115 (Token: FfoMUawOA9jI) and PXD054116 (Token: c1qB6Cwm8kOw) are related to Figure 4 and 5; PXD054100 (Token: hgMy360FLRFh) is related to Figure 7 and figure S9.

Microscopy data reported in this paper will be shared by the lead contact upon request.

This paper does not report original code.

Any additional information required to reanalyze the data reported in this paper is available from the lead contact upon request.

## Supplementary Materials

### Materials and Methods

#### Cell Culture

THP-1 (ATCC TIB-202) and mouse primary BMDM cells were cultured in Roswell Park Memorial Institute (RPMI, Gibco) 1640 with 10% FCS, and A549 cells were cultured in Dulbecco’s Modified Eagle Medium (DMEM, Gibco) supplemented with 10% FCS and Penicillin-Streptomycin (Gibco). Cells were maintained at 37°C with 5% CO_2_ before and after the treatment.

#### Antibodies and reagents

The antibodies and staining reagents used in western blot, immunofluorescence experiments and FACS sorting were: RIG-I Antibody (Abcam# ab180675), Ub Monoclonal Antibody (Cytoskeleton# # AUB01, clone:P4D1), α-tubulin Antibody (Merck#T9026), Myosin IIb Antibody (CST#3404), Actin Monoclonal Antibody (Sigma#ACTN05, clone:C4), COA7 Polyclonal Antibody (Invitrogen#COA7 Polyclonal Antibody), G3BP1 Polyclonal Antibody (Aviva#ARP37713_T100), EIF4H Monoclonal Antibody (CST#3469, clone D85F2), Goat anti-Mouse/Rabbit IgG (H+L) Superclonal^TM^ Recombinant Secondary Antibody Alexa Fluor 488/568 (Invitrogen#A28175 or A11031), CD45 (BD Bioscience#BV786), Ly6G (BD Bioscience#BV711), Ly6C (BD Bioscience#PE-cy7), CD11b (BD Bioscience#PerCP-Cy5.5), Mouse BD Fc Block™(BD Bioscience#553141), LIVE/DEAD™ Fixable Orange (602) Viability Kit (Thermo Fisher# L34983), and DAPI (Thermo Fisher# D1306).

#### Poly (I:C) treatment and virus infection

For poly (I:C) treatment, THP-1 cells and A549 cells were cultured as described and seeded at a density of 1x10^6^ cells per well in a 6-well plate with 2 mL medium. The cells were then treated with Poly (I:C) (2.5 μg/mL, with Xfect transfection reagent added) following the instructions. The details are outlined in figures (Fig. 1 to 7 and fig. S1 to 3, S9).

For virus infection, a stocked IAV X31 strain was used. The original titer was 4.4 x10^9^ PFU/mL. Cells were seeded 6∼8 hours before the infection. Cells were washed with EMEM infection medium [EMEM (ATCC), 10mM HEPES (Gibco), 0.125% BSA (Gibco), 0.5 μg/mL TPCK trypsin] and infected with MOI indicated in the corresponding figures. The zero-hour timepoint was harvested after 15 min at 37°C. Viruses were diluted in the infection medium.

#### Immunoblots

After Poly (I:C) treatment or virus infection, cells in suspension were harvested by centrifugation for 3 min (1,500 × g), while adherent cells were first digested with non-enzymatic dissociation reagent (Corning# 15313661) and then harvested by centrifugation. The supernatant was discarded, and cell pellets were lysed in Co-immunoprecipitation (Co-IP) buffer (10 mM Tris pH = 7.5, 100 mM NaCl, 0.5% NP-40, 0.5 mM EDTA) or RIPA buffer supplied with cOmplete^TM^ EDTA-free protease inhibitor cocktail (Roche#CO-RO) on ice for 30 min. The lysate was then centrifuged for 10 min at 4°C at 14,000 rpm, and the supernatant was transferred to a new 1.5 mL EP tube. Protein concentration in supernatant was determined by the Bradford assay. Equal amounts of proteins were loaded onto NuPAGE 4-12% gradient Bis-Tris gels (200V, 36 min), followed by membrane transfer using the Bio-Rad Turbo-blot transfer system. The Trans-Blot Turbo Mini 0.2 µm PVDF Transfer Packs (BIO-RAD# #1704156) was used for the membrane transfer process, and a mixed molecular weight program was chosen. After the membrane transfer, PVDF membranes were washed in miliQ water for 3 min, and then blocked in blocking buffer (5% skim milk, 20 mM Tris pH 7.5, 100 mM NaCl) at room temperature for 1 hr. Primary antibodies were then added in 5 mL blocking buffer and incubated with PVDF membranes at 4°C overnight. Then PVDF membranes were washed with TBS-T for 3 times, and then incubated in the appropriately diluted secondary antibody (in 1X TBS-T and including 5% skim milk) for 1 hr at room temperature with gentle rocking. Then the membrane was washed with TBS-T 3 times again. For detection, the membrane was incubated with Amersham ECL Detection Reagents (Cytiva# RPN3004) at room temperature for 3 min.

#### BMDM isolation and culture

Mice (C57BL/6, 8∼14 weeks old) were originally obtained from Charles River Company and were housed in specific-pathogen-free facilities at the Friedrich Miescher Institute for Biomedical Research (FMI). Mice were housed in temperature-controlled rooms on a constant 12-hr light-dark cycle with ad libitum access to water and food. The mice were euthanized by CO_2_ inhalation. Both legs of the sacrificed mice were sterilized in 70% ethanol. The attached tissues were removed by dissection with scissors and the isolated bones were crushed twice in a mortar with 5 mL RPMI 1640 medium. The suspension from the crushed bones was filtered through 100 μm cell strainer (Falcon#352360) to remove the debris. Filtered cells were centrifuged for 15 min at 500 × g at room temperature. The supernatant was removed, and cell pellets were resuspended in RPMI 1640 supplied with M-CSF-1 (100 ng/mL). The cell suspension from each leg was divided into four Nunc Square 25-cm dishes (Thermo Scientific#166508), with each plate containing 10 mL cell suspension. On the Day 3 post-seeding, 5 mL fresh RPMI 1640 medium supplied with M-CSF-1 was added to each plate. On Day 6, BMDMs were reseeded for new experiments or collected for the frozen storage. All animal experiments were approved by the FMI Animal Committee and the local veterinary authorities (Kantonales Veterinäramt of Kanton Basel-Stadt) and performed in accordance with the Swiss federal guidelines for animal experimentation.

#### TUBE pulldown

For a typical pulldown experiment, 60 μg C-terminal Biotinylated TUBE (prepared by following method in (*1*)) was pre-incubated with 15 uL streptavidin beads in a cold room for 30 min. The streptavidin-TUBE beads were then centrifuged to remove the unbound proteins in the supernatant and washed twice with the cell lysis buffer (Co-IP buffer:10 mM Tris pH = 7.5, 100 mM NaCl, 0.5% NP-40, 0.5 mM EDTA). Subsequently, the beads were incubated with 200 μg (total protein amount) cell lysate in the cold room and kept overnight on a rotator (10 rpm). Next day, the beads were washed by lysis buffer twice, followed by a TBS (10 mM Tris pH = 7.5, 150 mM NaCl) buffer wash. Captured protein was either analysed by immunoblot or by mass spectrometry. For immunoblot, the bead samples were dissolved in LDS sample buffer (ThermoFisher#NP0007) supplemented with 10 mM DTT and boiled for 5 min at 85°C before loading on 4%–12% Bis-Tris NuPAGE gels. Beads were spun down by centrifuge (12,000 x g, 1 min).

#### Immunofluorescence and microscopy

0.05 million cells were seeded in a 4-well chamber slide (ThermoFisher #154526). Following overnight culture, cells were treated or infected as depicted in the figures (Fig. 6 to 8 and fig. S9). Subsequently, the medium was aspirated, and cells were washed twice with ice-cold PBS. Cells were then fixed with 4% PFA for 10 min, permeabilized with 0.5% Triton X-100/PBS for 15 min, and blocked with blocking buffer (5% Goat serum in 0.1% Triton X-100/PBS) for 1 hr. Primary antibodies were applied to cells at a 1:150 dilution and incubated overnight at 4°C. The next day, cells were washed three times with PBS and incubated with fluorescent secondary antibodies in PBS or blocking buffer (1:250 dilution) for 1 hr at room temperature. DAPI (ThermoFisher#D1306) and Actin staining with Phalloidin 488 (CytoSkeleton #PHDG1) were diluted in PBS according to the product instructions and added to cells for 20 min at room temperature. Following three washes with PBS, cells were sealed with ProLong™ Diamond Antifade Mountant (ThermoFisher #P36965). Slides were inspected with spinning disk confocal microscope unit Yokogawa CSU W1 with Dual T2.

#### Labe-free mass spectrometry after TUBE pulldown in Poly (I:C) treated THP-1 cells

For label-free mass spectrometry (Figure 1 D), every replicate (n = 3) of the samples contained 10 million THP-1 cells. Cells were treated first with DMSO, TAK-243 (1 μM), Z-VAD-FMK (5 μM) and Bafilomycin A1 (500 nM) for 30 min, then Poly (I:C) (2.5 μg/mL) was transfected via X-fect transfecting reagent. Cells were harvested by centrifugation (1500 × g, 3 min) and lysed in Co-IP buffer. The cell lysate was incubated with streptavidin-TUBE beads following the method in “TUBE pulldown” section.

To digest the proteins on the beads, we used 10 μL Lys-C (0.2 mg/mL in 50 mM Hepes pH 8.5) and 50 μL digestion buffer (3 M GuaHCl, 20 mM EPPS pH 8.5, 10 mM CAA, 5 mM TCEP) to make master mix, and added 6 μL of this mixture to the beads from each 10 cm dish. After a short spin down (< 1000 × g) and 4 h incubation at 37°C, we added 17 μL 50 mM HEPES pH 8.5 as well as 1 μL 0.2 mg/mL trypsin to further digest the proteins at 37°C overnight. Next morning, another 1 μL trypsin was added to the solution for 6 hours digestion and quenched by adding 20% TFA in water to the final concentration of 1% TFA. Samples were analyzed by liquid chromatography–mass spectrometry (LC–MS)/MS on an EASY-nLC 1000 (Thermo Scientific) with a two-column set-up. The peptides were applied onto a peptide µPACᵀᴹ trapping column in 0.1% formic acid, 2% acetonitrile in H_2_O at a constant flow rate of 5 µl /min. Using a flow rate of 500 nl/min, peptides were separated at room temperature (RT°) with a linear gradient of 3 %–6 % buffer B in buffer A within 4 min followed by an linear increase from 6 to 22% in 55 min, 22 %–40 % in 4 min, 40%–80% in 1 min, and the column was finally washed for 10 min at 80% buffer B in buffer A (buffer A: 0.1% formic acid; buffer B: 0.1% formic acid in acetonitrile) on a 50 cm µPACᵀᴹ column (PharmaFluidics) mounted on a EASY-Spray™ source (Thermo Scientific) connected to an Orbitrap Fusion LUMOS (Thermo Scientific). The data were acquired using 120,000 resolutions for the peptide measurements in the Orbitrap and a top T (3 s) method with HCD fragmentation for each precursor and fragment measurement in the ion trap according to the recommendation of the manufacturer (Thermo Scientific). For the analysis, protein identification and relative quantification of the proteins was performed with MaxQuant v.1.5.3.8 using Andromeda as search engine, and label-free quantification (LFQ). The *H. sapiens* subset of the UniProt v.20190410 combined with the contaminant database from MaxQuant (*2*) was searched and the protein and peptide FDR were set to 0.01. The LFQ intensities estimated by MaxQuant were analyzed with the einprot R package (https://github.com/fmicompbio/einprot) v0.3.1. Features classified by MaxQuant as potential contaminants or reverse (decoy) hits or identified only by site, as well as features identified based on a single peptide or with a score below 10, were filtered out. The LFQ intensities were log_2_ transformed and missing values were imputed using the ‘MinProb’ method from the imputeLCMD R package v2.0 with default settings. Pairwise comparisons were performed using limma v3.50.0, considering only features with at least 2 non-imputed values across all the samples in the comparison. Estimated log_2_-fold changes and P-values (moderated t-test) from limma were used to construct volcano plots (*3*).

#### TUBE-TMT mass spectrometry analysis of Poly (I:C) induced ubiquitinome dynamics

THP-1 cells were treated with 2.5 μg/mL Poly (I:C) and harvested triplicate biological samples at 0 hr, 3 hr, 6 hr, 12 hr, and 24 hr post Poly (I:C) treatment. For the 0 hr timepoint, treating the cells with X-fect transfection reagent alone was used as a blank control. Cell collection was carried out via centrifugation at 1,500 x g for 3 min, followed by lysis using Co-IP buffer. Subsequently, a “TUBE pulldown” method was employed on the cell lysate obtained from each replicate containing 10 million cells. After the TUBE pulldown, the samples were removed from the beads using a magnetic rack. 0.25 mg of TMT reagent dissolved in 20 uL anhydrous acetonitrile were added to the supernatant. After 1 hr, labelling reaction was quenched using hydroxylamine, samples were pooled and fractionated by off-line high pH fractionation. The fractionation was carried out on a YMC Triart C18 0.5 × 250 mm column (YMC Europe GmbH) using the Agilent 1100 system (Agilent Technologies). A total of 96 fractions were collected for each experiment and concatenated into 48 fractions as previously described (*4*). Samples were analyzed by LC–MS/MS on an EASY-nLC 1000 (Thermo Scientific) with a column set-up. The peptides were applied onto a peptide µPACᵀᴹ column in 0.1% formic acid, 2% acetonitrile in H_2_O at a constant flow rate of 5 µL/min. Using a flow rate of 500 nL/min, peptides were separated at RT with a linear gradient of 3%–6% buffer B in buffer A in 8 min followed by an linear increase from 6 to 22% in 60 min, 22%–40% in 22 min, 40%–80% in 1 min, and the column was finally washed for 9 min at 80% buffer B in buffer A (buffer A: 0.1% formic acid; buffer B: 0.1% formic acid in acetonitrile) on a 50 cm µPACᵀᴹ column (PharmaFluidics) mounted on a EASY-Spray™ source (Thermo Scientific) connected to an Orbitrap Fusion LUMOS (Thermo Scientific). The data were acquired using 120,000 resolution for the peptide measurements in the Orbitrap and a top T (3 s) method with HCD fragmentation for each precursor and fragment measurement in the orbitrap at 50,000 resolution according the recommendation of the manufacturer (Thermo Scientific). Thermo RAW files were processed using Proteome Discoverer 2.5 software (Thermo Fisher) as described in the manufacturer’s instructions. Briefly, the Sequest search engine was used to search the MS2 spectra against the *Homo sapiens* UniProt database (downloaded on 03/05/2021) supplemented with common contaminating proteins. For total proteome analysis, cysteine carbamidomethylation and TMT tags on lysine and peptide N-termini were set as static modifications, whereas oxidation of methionine residues and acetylation protein N-termini were set as variable modifications. The TMT intensities estimated by PD were analyzed with the einprot R package (https://github.com/fmicompbio/einprot) v0.4.1. Features classified by PD as potential contaminants or reverse (decoy) hits, as well as features identified based on a single peptide were filtered out. The TMT intensities were log_2_ transformed and missing values were imputed using the ‘MinProb’ method from the imputeLCMD R package v2.0 with default settings. Pairwise comparisons were performed using limma v3.50.0, considering only features with at least 2 non-imputed values across all the samples in the comparison. Estimated log_2_-fold changes and P-values (moderated t-test) from limma were used to construct volcano plots (*3*).

#### diGly-TMT mass spectrometry analysis of ubiquitinome in Poly (I:C) treated THP-1 cells

5 million THP-1 cells were seeded into a T75 flask containing 10 mL of fresh RPMI1640 medium and subsequently treated with Poly (I:C) at a final concentration of 2.5 μg/mL, as outlined in the experimental procedure (Fig. 3A). Three timepoints were selected: 0 hr, 6 hr, and 24 hr, with each timepoint represented by 12 flasks (3 biological replicates, each replicate had 4 flasks). Cells were harvested by centrifugation at 1,500 x g for 3 min and lysed using 1% SDC lysis buffer. A portion of the cell lysate was utilized for the total protein quantification, while the remainder was subjected to K-GG motif enrichment following the protocol provided by the PTMScan® HS Ubiquitin/SUMO Remnant Motif (K-ε-GG) Kit (Cell Signaling#59322). The samples for total protein quantification and antibody enriched peptides were labelled with TMT reagent, pooled and fractionated by off-line high pH fractionation. The fractionation was carried out on a YMC Triart C18 0.5 × 250 mm column (YMC Europe GmbH) using the Agilent 1100 system (Agilent Technologies). A total of 96 fractions were collected for each experiment and concatenated into 48 fractions as previously described (*4*). Samples were analyzed by LC–MS/MS on an EASY-nLC 1000 (Thermo Scientific) with a column set-up. The peptides were applied onto a peptide µPACᵀᴹ column in 0.1% formic acid, 2% acetonitrile in H_2_O at a constant flow rate of 5 µL/min. Using a flow rate of 500 nL/min, peptides were separated at RT with a linear gradient of 3%–6% buffer B in buffer A in 8 min followed by an linear increase from 6 to 22% in 65 min, 22%–40% in 25 min, 40%–80% in 1 min, and the column was finally washed for 9 min at 80% buffer B in buffer A (buffer A: 0.1% formic acid; buffer B: 0.1% formic acid in acetonitrile) on a 50 cm µPACᵀᴹ column (PharmaFluidics) mounted on a EASY-Spray™ source (Thermo Scientific) connected to an Orbitrap Fusion LUMOS (Thermo Scientific). The data were acquired using 120,000 resolution for the peptide measurements in the Orbitrap and a top T (3 s) method with HCD fragmentation for each precursor and fragment measurement in the orbitrap at 50,000 resolution according the recommendation of the manufacturer (Thermo Scientific). Thermo RAW files were processed using Proteome Discoverer 2.5 software (Thermo Fisher) as described in the manufacturer’s instructions. Briefly, the Sequest search engine was used to search the MS2 spectra against the *Homo sapiens* UniProt database (downloaded on 03/05/2021) supplemented with common contaminating proteins. For total proteome analysis, cysteine carbamidomethylation and TMT tags on lysine and peptide N-termini were set as static modifications, whereas oxidation of methionine residues and acetylation protein N-termini were set as variable modifications. For modified peptides analysis, TMT tags on lysine were set as a variable modification and in addition TMT6plexDiGly / +343.206 Da (K) modification was included. The TMT intensities estimated by PD were analyzed with the einprot R package (https://github.com/fmicompbio/einprot) v0.5.13. Features classified by PD as potential contaminants or reverse (decoy) hits, as well as features identified based on a single peptide were filtered out. The TMT intensities were log_2_ transformed and missing values were imputed using the ‘MinProb’ method from the imputeLCMD R package v2.0 with default settings. Pairwise comparisons were performed using limma v3.50.0, considering only features with at least 2 non-imputed values across all the samples in the comparison. Estimated log_2_-fold changes and P-values (moderated t-test) from limma were used to construct volcano plots (*3*).

#### diGly-TMT mass spectrometry analysis of ubiquitinome in IAV infected THP-1 cells

For IAV infection, 5 million THP-1 cells were seeded into a T75 flask containing 10 mL of fresh RPMI1640 medium. And IAV strain X31 was used for infection (MOI = 10). The details of infection procedures and medium were described above. At 24 hr and 48 hr post-infection, cells were collected by centrifuge (1,500 x g, 3 min). Part of cells were used for RNA extraction with single cell RNA extraction kit (Norgen#SKU 51800). Extracted RNA was sent for total RNA sequencing. The rest of cells were used for diGly peptide enrichment followed by TMT-based quantification and total protein abundance analysis with TMT-based quantification. Cells were dissolved in 200 uL or 400 uL 1% SDC 50 mM HEPES pH = 8.5 sonicated 10 sec at max power using the tip sonicate, supplemented with 2mM TCEP and 10 mM CAA for 30 min and trypsin 1:100, Lys-C 1:500. The samples were incubated overnight at 37°C. In the morning Trypsin was added at a 1:100 dilution. The samples were labelled with TMTpro reagent and fractionated by off-line high pH fractionation. The fractionation was carried out on a YMC Triart C18 0.5 × 250 mm column (YMC Europe GmbH) using the Agilent 1100 system (Agilent Technologies). A total of 96 fractions were collected for each experiment and concatenated into 48 fractions as previously described (*4*). Samples were analyzed by LC–MS/MS on a Van Vanquish Neo UHPLC system (Thermo Scientific) with a two-column set-up. The peptides were applied onto a PepMap Neo trap (Thermo Fisher) in 0.1 % formic acid, 2% acetonitrile in H_2_O. Using a flow rate of 200 nL/min, peptides were separated at RT with a linear gradient of 2%–7% buffer B in buffer A in 2.5 min followed by an linear increase from 7 to 20% in 65 min, 20%–30% in 22 min, 30%–36% in 8 min, 36%–45% in 3 min, 45%–100% in 2 min, and the column was finally washed for 5 min at 100% buffer B in buffer A (buffer A: 0.1% formic acid; buffer B: 0.1% formic acid in 80% acetonitrile) on 15-cm EASY-Spray^™^ C18 column (ES75150PN, Thermo Fisher) (PharmaFluidics) mounted on a EASY-Spray™ source (Thermo Scientific) connected to an Orbitrap Fusion Eclipse (Thermo Scientific). The data for modified peptide measurements were acquired using 120,000 resolutions for the peptide measurements in the Orbitrap and a top T (3 s) method with HCD fragmentation for each precursor with charge 3+ and higher and fragment measurement in the orbitrap at 50,000 resolution according the recommendation of the manufacturer (Thermo Scientific). For proteome measurements the FAIMS device was employed with -40, -60, -80 compensation voltages in combination with real time search against the *Homo sapiens* UniProt database (downloaded on 05/2021). Thermo RAW files were processed using Proteome Discoverer 2.5 software (Thermo Fisher) as described in the manufacturer’s instructions. Briefly, the Sequest search engine was used to search the MS2 spectra against the *Homo sapiens* UniProt database (downloaded on 03/05/2021) supplemented with common contaminating proteins. For total proteome analysis, cysteine carbamidomethylation and TMT tags on lysine and peptide N-termini were set as static modifications, whereas oxidation of methionine residues and acetylation protein N-termini were set as variable modifications. For modified peptides analysis, TMT tags on lysine were set as a variable modification and in addition TMTproDiGly /+418.250 Da (K) modification was included. The TMT intensities estimated by PD were analyzed with the einprot R package (https://github.com/fmicompbio/einprot) v0.7.3. Features classified by PD as potential contaminants or reverse (decoy) hits, as well as features identified based on a single peptide were filtered out. The TMT intensities were log_2_ transformed and missing values were imputed using the ‘MinProb’ method from the imputeLCMD R package v2.0 with default settings. Pairwise comparisons were performed using limma v3.50.0, considering only features with at least 2 non-imputed values across all the samples in the comparison. Estimated log_2_-fold changes and P-values (moderated t-test) from limma were used to construct volcano plots (*3*).

#### RNA sequencing of IAV infected THP-1 cells

Total RNA of 10% of the IAV infected THP-1 cells (from previous section “diGly-TMT mass spectrometry analysis of ubiquitinome in IAV infected THP-1 cells”) was extracted with the the Single Cell RNA Purification Kit (Norgen#SKU 51800). The samples were further processed at the FMI genomics Facility. The results were analyzed with Galaxy platform (*5*). The RNA sequencing reads were firstly aligned to reference genome (GRCh38.primary_assembly) and then quantified by QuasR (*6*). Original data and RNA sequencing details were deposited at ArrayExpress (accession E-MTAB-14283).

#### diGly-TMT mass spectrometry analysis of ubiquitinated proteins in primary monocytes from IAV infected mice

A total of 30 mice aged 8 to 10 weeks were utilized in the study and divided into three groups: 10 mice for non-infected controls, 10 mice infected with IAV (PR8, 200 PFU) and sacrificed on day 3 post-infection, and 10 mice infected and sacrificed on day 7 post-infection. All animal experiments were approved by the FMI Animal Committee and the local veterinary authorities (Kantonales Veterinäramt of Kanton Basel-Stadt) and performed in accordance with the Swiss federal guidelines for animal experimentation under the license 2825. Following the sacrifice, lung tissues were collected in PBS with 2% FCS and minced into homogeneous pastes (∼1 mm in size) using scissors. Minced tissues from two mice were then incubated with 12 mL dissociation medium (75% RPMI1640 + 10% Collagenase/Hyaluronidase + 15% Dnase I solution, 1 mg/mL) at 37°C for 40 min. The tissues were subsequently pushed through a 70 μm nylon mesh strainer using the rubber end of a syringe plunger. The strainer was washed with 2 mL PBS (+2% FCS), and the filtered solution was centrifuged (300 x g, 10 min) to pellet cells, with the supernatant carefully removed. The pellet was then incubated with 20 mL ammonium chloride at room temperature for 10 min, followed by filling to 50 mL with RPMI and centrifugation (300 x g, 10 min). The resulting pellet was resuspended, and cell quantification performed in 5 mL PBS (+ 2% FCS).

For antibody staining, 2 mL diluted Fc blocker (prepared by adding 300 μL stock solution to 10 mL PBS + 2% FCS) was added to resuspended cells for a 20-minute incubation on ice. Cells were then centrifuged (500 x g, 5 min) at 4°C, and the supernatant was discarded. Subsequently, 10 mL antibody mixer (PBS containing 10 μL Live/Dead staining and 10 μL antibodies for CD45, CD11b, Ly6G, and Ly6C, stock concentration: 0.2 mg/mL) was added to the cells and incubated together on ice for 30 to 40 min. The cells were washed with FACS buffer (PBS + 2% FCS) by centrifugation twice, then resuspended in 2 mL FACS buffer and filtered with a 30 μm cell strainer. Finally, cells were sorted using a Sony Sorter MA900. For mass spectrometry analysis, experimental design was as shown in figure S6C.

Each sample was lysed with 40 uL lysis buffer (1% SDC pH = 8, 50 mM HEPES, 10 mM CAA, 2 mM TCEP). 0.5 μg Lys-C and 2 μg of trypsin were added to the pellets and incubated at 37°C overnight. Later samples were sonicated in the morning using Bioruptor (6 cycles, 30 sec on and 15 sec off), and 380 units Benzonase added and incubated for 15 min. Further analysis was caried out as described in the section: “*diGly-TMT mass spectrometry analysis of ubiquitinome in IAV infected THP-1 cells.”*

#### Lentivirus cell line establishment

Lentivirus transfer plasmids (original vector was bought from Addgene, #19070) (*7*) expressing GFP, GFP-COA7-NLS, GFP-COA7-NES, and mCherry-EIF4H-IRES-GFP, were obtained from TWIST Bioscience. 10 μg transfer plasmids were co-transfected with 1 μg packaging plasmids (expressing Gag, Pol, Rev, and Tat) into 1.6 million 293T cells in a 10 cm dish using 30 μL FuGENE HD transfecting reagent. After 3 days of culturing, the supernatant containing lentivirus was collected and precipitated using LentiX concentrator (Takara#631231) according to the manufacturer’s instructions. The virus pellet was resuspended in complete medium and added to target cells. Following 3 days of lentivirus infection, a portion of cells was frozen as a “cell pool,” while the remaining cells were sorted by fluorescent signal with FACS into 96-well plates, with each well containing a single cell. After 2 to 4 weeks of culturing, single-cell colonies were transferred individually to 12-well plates and cultured for an additional week. Cells from the 12-well plates were then subjected to immunoblot analysis to confirm the expression of the target protein. Correctly expressing single-cell colonies were frozen at -80°C and labelled according to their original positions on the 96-well plate. One day later, the cells were transferred to liquid nitrogen tank for long-term storage.

#### Identification of COA7 interacting proteins by affinity purification - mass spectrometry analysis

Treated GFP-COA7 A549 cells were harvested with ice-cold PBS from a 10 cm dish with Cell Lifter (Corning#3008) and centrifuged (1,500 x g, 3 min) at 4°C. The pelleted samples were rapidly frozen at -20°C. The frozen pellet was lysed as described previously. The lysed cell extracts were subjected to high-speed centrifugation (13,000 x g for 10 min, 4°C) to remove the cell debris. GFP-tagged COA7 were pulled down with GFP-Trap Magnetic agarose beads (Chromotek#gtm-20) equilibrated with the same lysis buffer. Lysates were incubated with GFP-Trap beads for 30∼60 min in a cold room, then beads were spun down at 1,000 x g for 2 min. The beads were washed with lysis buffer twice, and finally washed with (10 mM Tris, pH 7.5, 150 mM NaCl) buffer twice. The bead samples were proceeded with the on-beads digestion protocol (same as “*Label-free mass spectrometry after TUBE pulldown in Poly (I:C) treated THP-1 cells*” section) for label-free mass spectrometry analysis (see above) at the FMI proteomic facility.

#### Analysis of EIF4H degradation by immunoblot and FACS

The mCherry-EIF4H-IRES-GFP A549 cell lines were established according to the previous description (Lentiviral plasmid was synthesized by TWIST Bioscience). For Poly (I:C) treatment, 0.2 million cells were seeded into each well of a 6-well plate. After overnight incubation, cells were treated with 5 μg/mL (final concentration) Poly (I:C) along with other chemicals (e.g., Bortezomib) as indicated in the Figure 6. However, for IAV infection, cells were infected with IAV X31 at 6 to 8 hours post-seeding at the indicated MOI. Following 24 hours of treatment or infection, cells were detached using Trypsin-EDTA dissociation medium and centrifuged at room temperature (1,500 x g, 3 min). For FACS analysis, the cell pellet was resuspended in FACS buffer (2% FCS in PBS) and filtered through a 40-µm cell strainer (Corning #08-771-1). Fluorescent signals were recorded using a BD LSRII SORP Analyzer. For immunoblot analysis, the cell pellet was washed twice with PBS and then lysed using a previously described lysis buffer, following the procedures outlined in the “immunoblot” section.

#### Screening of E2s that are responsible for EIF4H ubiquitination

Horizon Discovery ON-Targetplus siRNAs targeting 39 host E2 enzyme genes were randomly distributed over three 96-well plates, with 3 distinct siRNAs targeting each gene. For reverse transfection, working plates were prepared with 20 nM siRNA in 15 µL serum-less DMEM, into which 0.15 µL of Lipofectamine RNAiMax (Invitrogen# 13778075) transfection reagent in 15 µL of serum-less DMEM was added and mixed thoroughly. Following a 45 min incubation at room temperature, 2,000 A549 cells in 70 µL of growth medium (DMEM, 10% FCS and NEAA) were added to each well. After 72 h incubation at 37°C, 5% CO_2_, the cells were washed once with growth medium and transfected with 0.5 µg of Poly (I:C) LMW (InvivoGen) per well using 0.15 µL per well of X-fect transfection reagent (Takara Bio# 631318).

Cells transfected with non-targeting siRNAs but without Poly (I:C) transfection was used as the negative control for EIF4H degradation. 12 hours after Poly (I:C) transfection, cells were fixed with 4% PFA in PBS, and stained for EIF4H using anti-EIF4H (D85F2) rabbit mAb (1:150 dilution) and for the nuclei with Hoechst dye (1:2000). The cell membrane was demarcated by staining with WGA-AF680 (1:1000). Automatic spinning-disk confocal image acquisition was carried out using the 20x objective lens of the Cell Voyager™ CQ1 Benchtop High-Content Analysis System (Yokogawa Electric Corporation). Z-Stack Summation (ZSUM) images obtained from 5 Z-stack images were used. Cell Profiler was used to perform image segmentation and to calculate mean intensities of EIF4H per individual cell. Data were processed using R to obtain mean intensity and mean cell number plots.

#### TCID_50_ quantification of IAV infectivity in A549 cells

IAV (X31, MOI = 0.1) infected WT A549, GFP-expressing A549, and EIF4H over-expressing A549 cells in infection medium by following the previous method in “virus infection”. Infection was duplicated, with each replicate containing two technical repeats. Supernatants were collected at 48 hr post-infection. For TCID50 quantification, 10,000 MDCK cells were seeded per well in 96 well plate. After overnight culturing in the incubator at 37°C, cell confluency should be > 90%. Then the supernatant with virus (original supernatant) was diluted 10 times in the infection medium. This diluted supernatant was further serially diluted 10-fold in infection medium, and 200 uL diluted medium was added to MDCK cells in 96 well plate. For example, the first row was 100 times diluted supernatant from the original one, and the eighth row was 10^10^ diluted supernatant from the original one.

After 3 days post-infection, the supernatants were removed, and cells were fixed with 50 µL 4% PFA for 10 min. Then 50 µL crystal violet solution (Sigma# HT901) was added to each well and incubated for 15 min. After the incubation, excess crystal violet solution was removed: the virus-infected cells detached while non-infected cells were intact and purple. The cytopathic effect was quantified accordingly.

#### RT-qPCR

For THP-1 cells, 1 million cells were seeded into individual wells of a 6-well plate containing 2 mL fresh culture medium. The treatment was administered immediately following seeding, in accordance with the experimental design indicated in the figures (Fig. 1, 6, 7 and fig. S1, S9). In the case of A549 cells, a reduced cell number (0.1 to 0.3 million cells) was seeded one night prior to the treatment. Subsequently, the cells were harvested, and the total RNA was extracted using the Single Cell RNA Purification Kit (Norgen# SKU 51800), followed by a reverse transcription using the ImProm-II™ Reverse Transcription System (Promega# A3800). The transcribed cDNA was then diluted sixfold using RNase/DNase-free water, and qPCR primers (sequences provided in table S10) were added at a final concentration of 0.5 μM. The qPCR reactions were performed using 2x FastStart Universal SYBR Green Master (Merck# 4913850001) and Applied Biosystems Real-Time PCR Instruments. All samples were run in triplicate to ensure the robustness of the results.

#### Statistical analysis

For all the mass spectrometry analyses, we considered the protein or peptide with Log_2_FC > 1 or < -1 and adj.p-value (corrected p-value by using the Benjamini-Hochberg method) < 0.01 as increased or decreased. Other statistical analysis details have been indicated in the figure legends.

To define the proteins with increased ubiquitination in Figure 4 and 5, we calculated the adj.p-value for proteins with multiple diGly peptides in THP-1 under IAV infection, and aggregated the peptide-level p-values for each protein using Simes method, adjusted for multiple testing across proteins using the Benjamini-Hochberg method, and then got the adj.p-value.

Data were plotted using Prism v.10.2.3 (GraphPad) and presented as the means of multiple experiments, with error bars showing the ± SD. The significance of results was determined using ordinary one-way ANOVA or unpaired T-tests, as indicated in figure legends, and p < 0.05 was considered as significant.

**Fig. S1.**
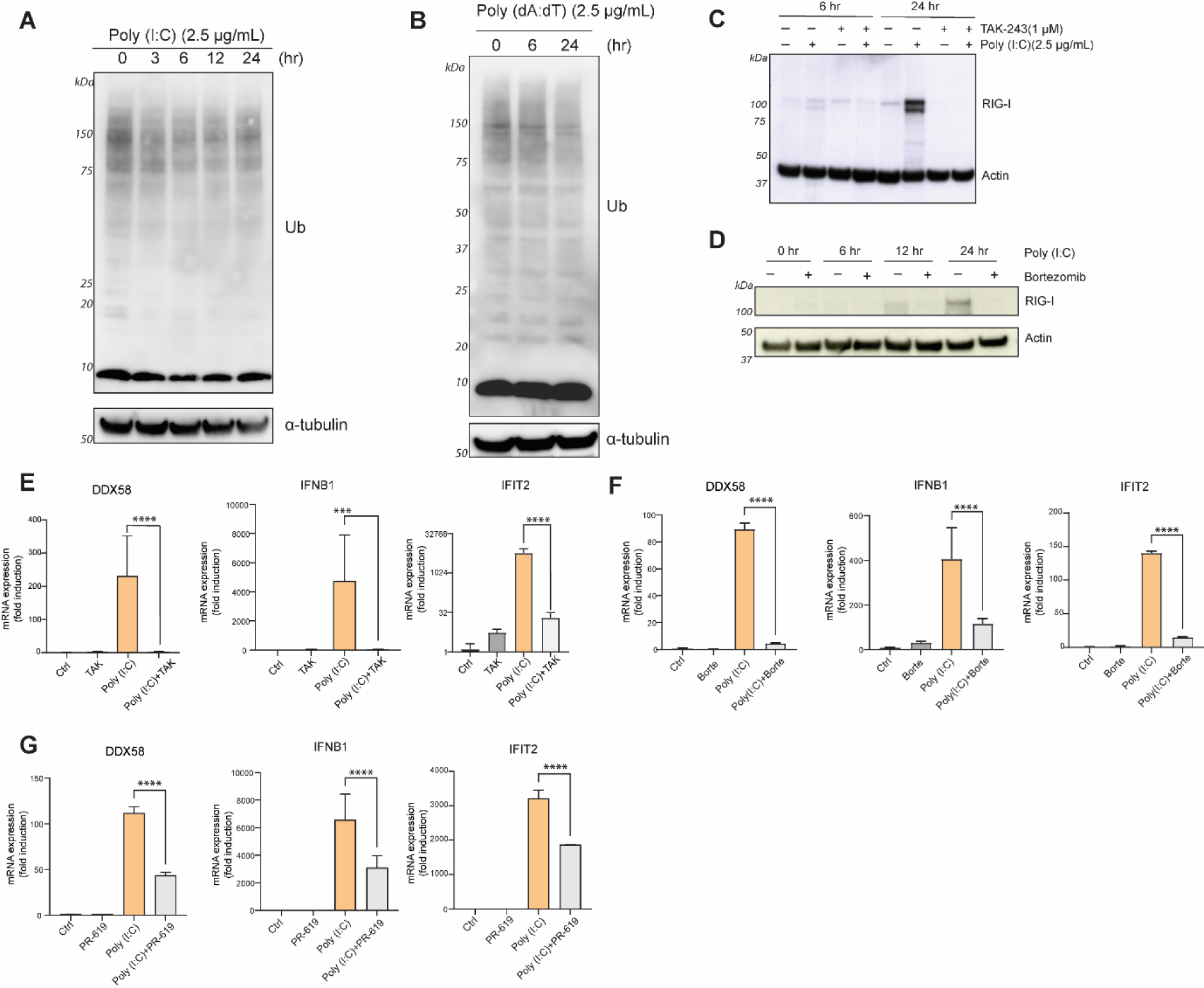
Interfering with ubiquitination inhibits proinflammatory and antiviral cytokines release. (**A**) Immunoblotting of global ubiquitination in Poly (I:C) treated A549 cells. Cells were treated as indicated, and α-tubulin served as internal control. (**B**) Immunoblotting of global ubiquitination in Poly (dA:dT) treated THP-1 cells. Cells were treated as indicated, and α-tubulin served as internal control. (**C**) Immunoblotting of TAK-243 effect on RIG-I activation. THP-1 cells were treated with 2.5 μg/mL Poly (I:C) for 6 hr or 24 hr; TAK-243 (1 μM) was added together with Poly (I:C) transfection. (**D**) Immunoblotting of Bortezomib effect on RIG-I activation. THP-1 cells were transfected with 2.5 μg/mL Poly (I:C) and incubated for the indicated time; Bortezomib (1 μM) was added together with Poly (I:C) treatment. (**E** to **G**) RT-qPCR result of TAK-243 (E), Bortezomib (F), and PR-619 (G) effect on Poly (I:C) treated THP-1 cells at 6 hr (n = 3). Treatments conditions were as above. The 2^–ΔΔCt^ method was used for relative mRNA quantification. GAPDH was the reference gene. Statistical analysis was performed with one-way ANOVA test. ***, p < 0.001; ****, p < 0.0001; ns, not significant

**Fig. S2.**
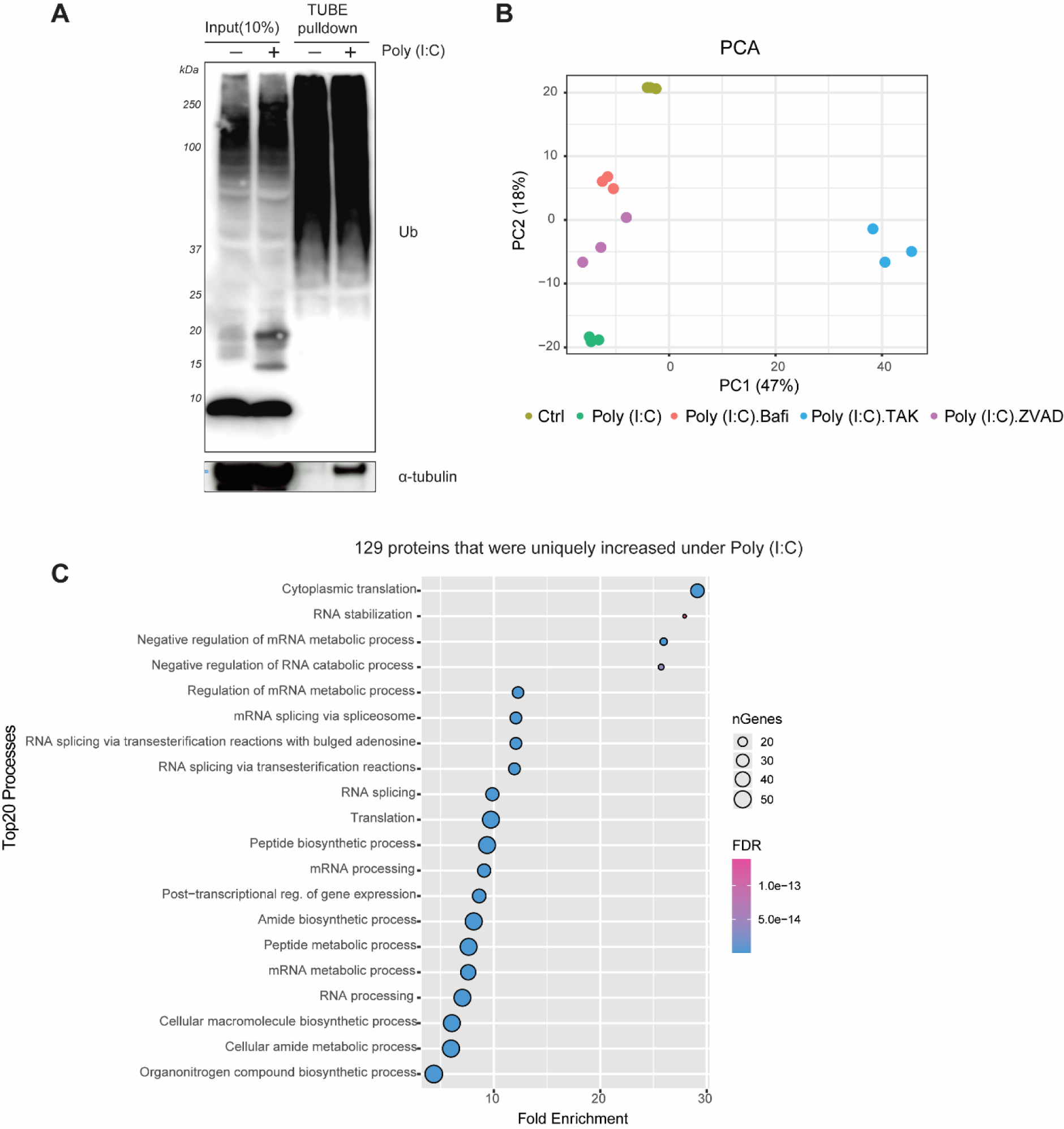
TUBE based ubiquitinated proteins enrichment. (**A**) TUBE efficiently captures ubiquitinated proteins from THP-1 cell lysates with or without Poly (I:C) treatment (2.5 μg/mL, 6 hr). (**B**) PCA analysis of the proteomic data from each treatment. As visible, TAK-243 treated cells were dramatically different from the other 4 conditions on the PC1 axis. (**C**) GO_BP analysis of 129 proteins that were uniquely increased with TUBE enrichment in Poly (I:C) treated THP-1 cells. TOP 20 processes were sorted by FDR (< 0.05) and organized by Fold Enrichment.

**Fig. S3.**
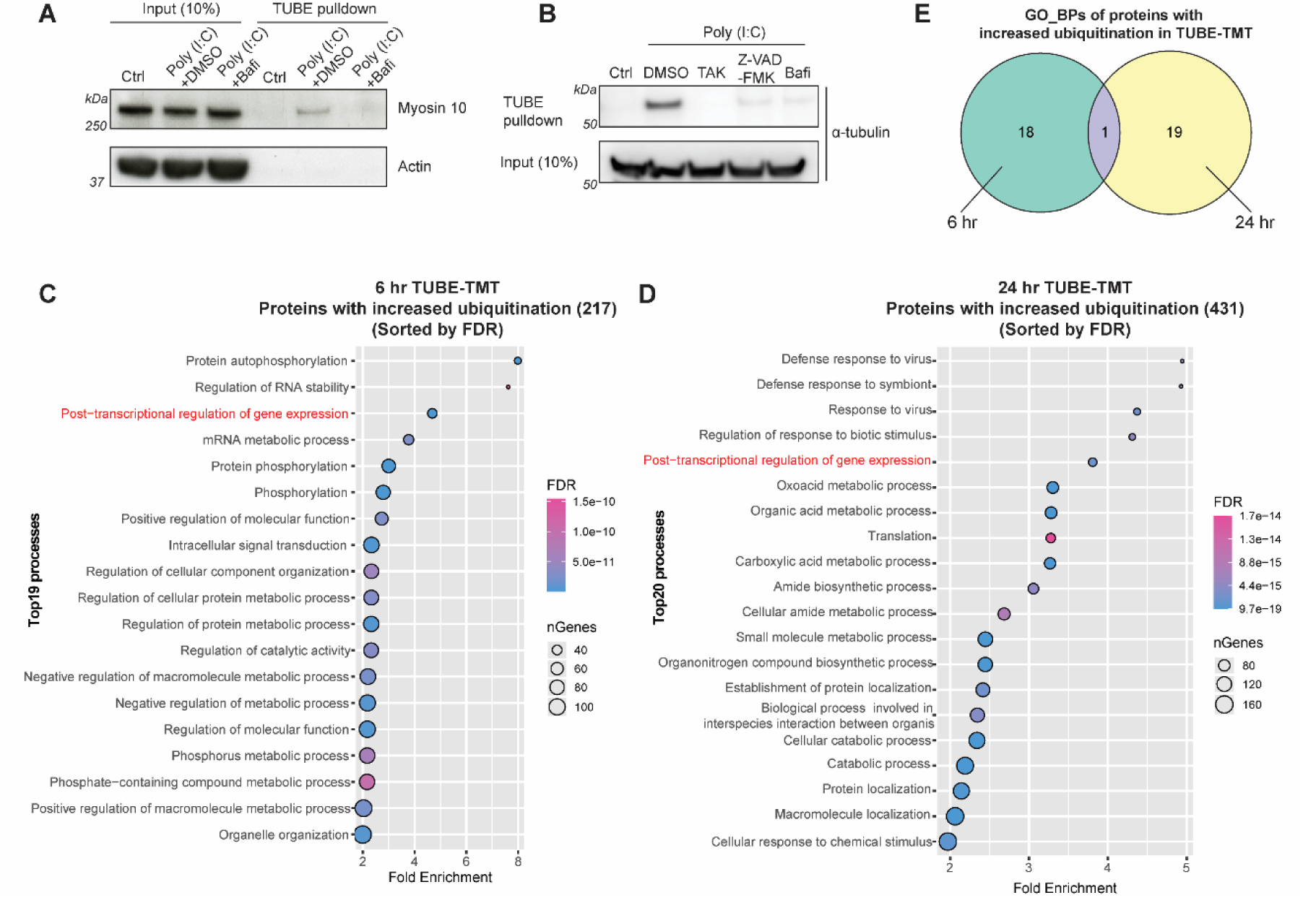
Ubiquitination of Myosin 10 and α-tubulin and analysis of TUBE-TMT proteomics. (**A**) Immunoblotting for Myosin 10 ubiquitination by TUBE pulldown with THP-1 cell lysates. Bafilomycin A1 (500 nM) was added together with Poly (I:C) transfection (2.5 μg/mL, 6 hr). (**B**) Immunoblotting for α-tubulin ubiquitination by TUBE pulldown with THP-1 cell lysates. Cells were transfected with Poly (I:C) (2.5 μg/mL) and were analyzed after 6 hr; TAK-243 (1 μM), Z-VAD-FMK (5 μM) and Bafilomycin A1 (500 nM) were added together with Poly (I:C) transfection. (**C, D**) GO_BP analysis of 6 hr and 24 hr proteins with increased ubiquitination (corresponding to Figure 2C and E with FC > 2, adj.p-value < 0.01) in TUBE-TMT proteomic analysis. TOP 20 processes were sorted by FDR (< 0.01) and organized by Fold Enrichment. For 6 hr, only 19 processes are presented, as the 20^th^ process had a low fold enrichment (< 2) and was excluded. Red colored process is common to both timepoints. (**E**) Venn diagram of the processes identified in (C) and (D).

**Fig. S4.**
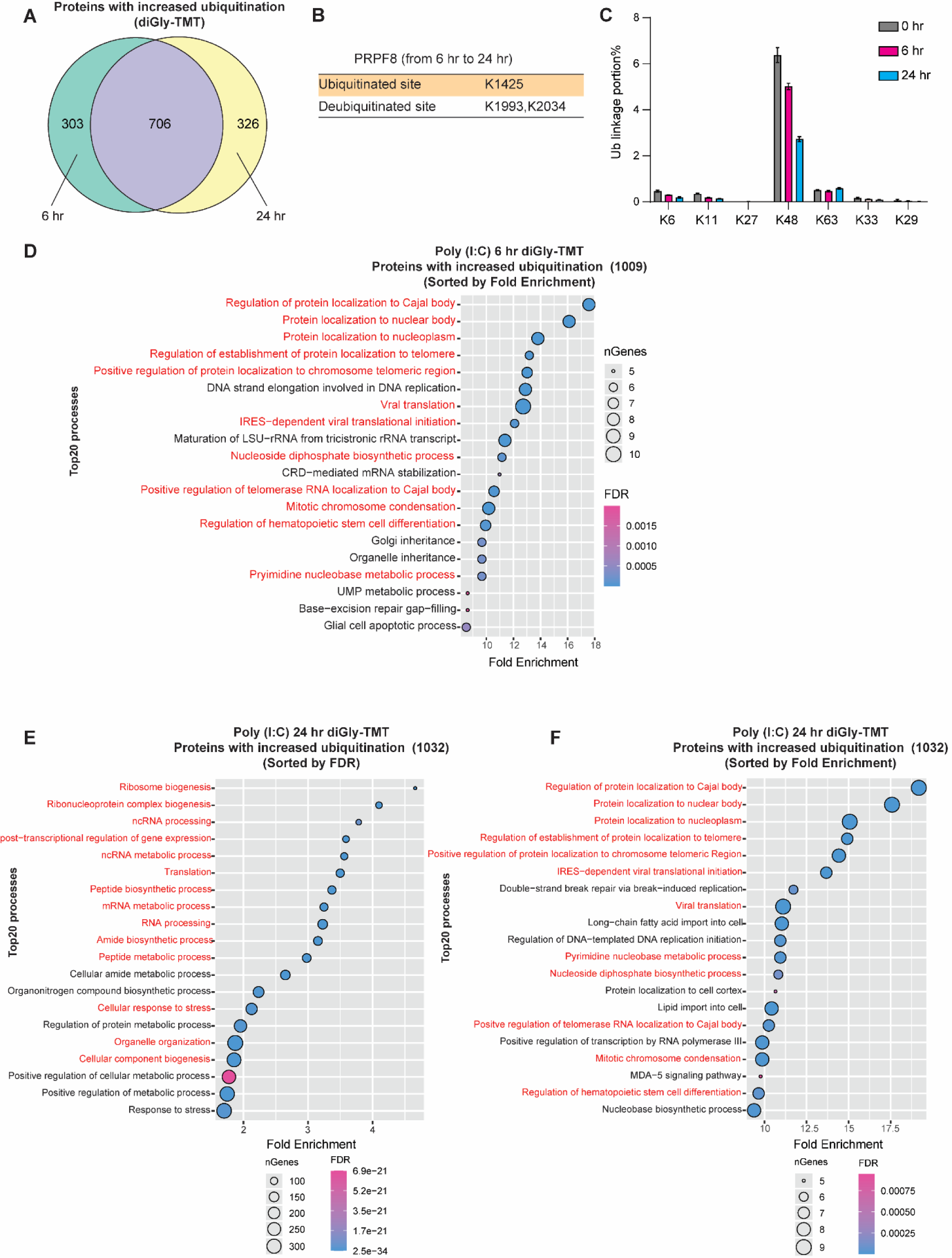
Comparison of Poly (I:C)-induced 6 hr and 24 hr increased ubiquitination in diGly-TMT mass spectrometry. (**A**) Venn diagram for proteins with increased ubiquitination (identified in diGly-TMT mass spectrometry analysis) at 6 hr or 24 hr post Poly (I:C) treatment. (B) PRPF8 ubiquitinated and deubiquitinated sites under Poly (I:C) treatment in THP-1 cells. (C) Portion of all the Ub chain linkages’ peptides in total diGly peptides (supplemental to Fig. 3J). (**D**) GO_BP analysis of proteins with increased ubiquitination (Poly (I:C) 6 hr diGly-TMT). Top 20 Biological processes were sorted and organized by Fold Enrichment (FDR < 0.01). The red color represents the processes that are common to both timepoints. (**E**) and (**F**) GO_BP analysis of proteins with increased ubiquitination (Poly (I:C) 24 hr diGly-TMT). Top 20 Biological processes were sorted by FDR and organized by Fold Enrichment (E) or sorted and organized by Fold Enrichment (FDR < 0.01) (F). The red color represents the processes that are common to both timepoints.

**Fig. S5.**
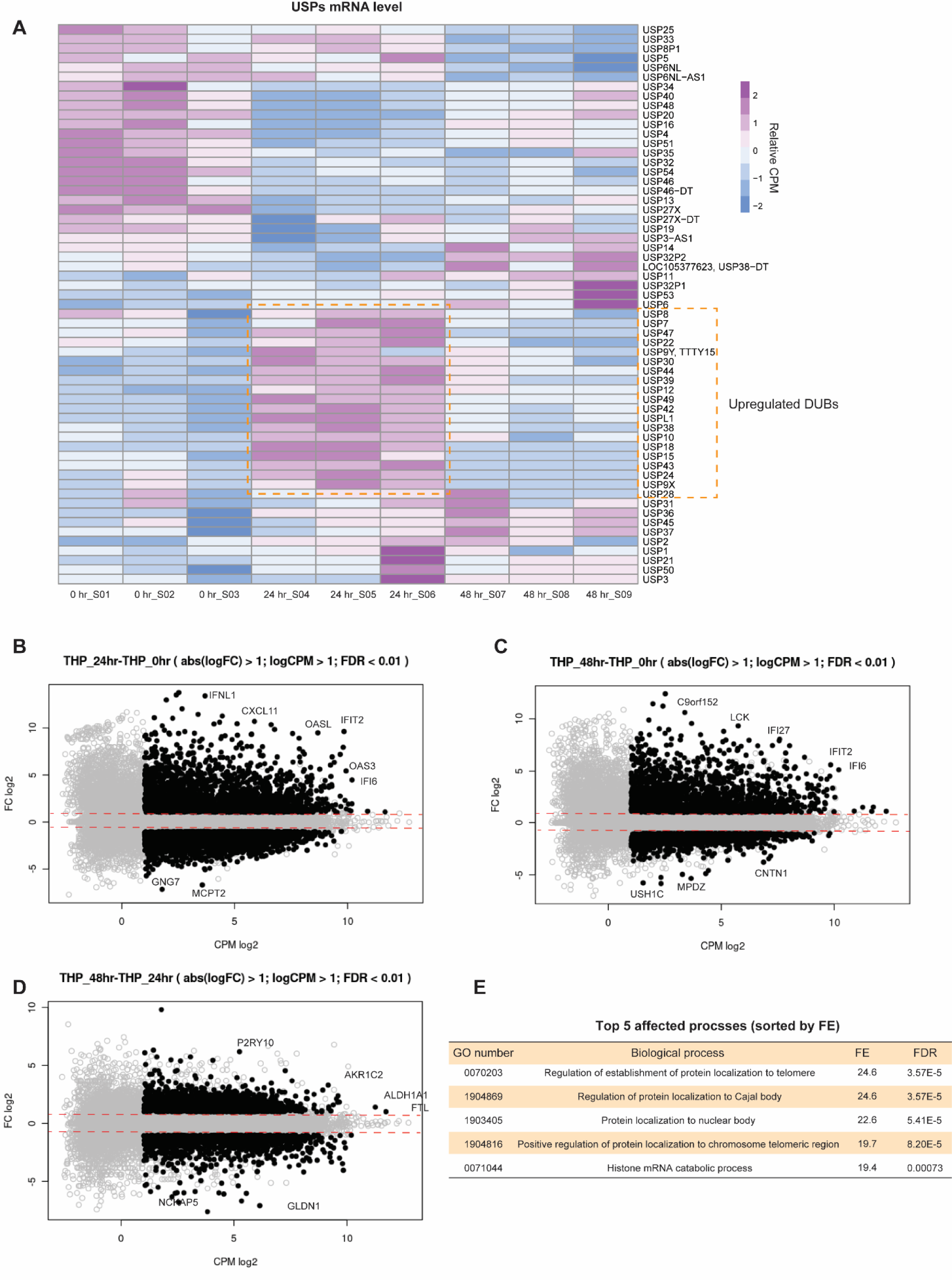
RNA sequencing of IAV infected THP-1 cells. (**A**) Heatmap of USPs RNA level in IAV infected THP-1 cells. S is short for sample. Orange dashed rectangle indicates IAV induced USPs’ expression at 24 hr post-infection. (**B** to **D**) MA plots of RNA sequencing for THP-1 cells. One dot represents a gene. (B) 24 hr vs 0 hr comparison, (C) 48 hr vs 0 hr and (D) 48 hr vs 24 hr in IAV infected THP-1 cells. Red dashed line represents the log_2_FC = ±1. (**E**) GO_BP analysis of proteins with increased ubiquitination (n = 453) at 24 hr post IAV infection. The top 5 biological processes were sorted and organized by Fold Enrichment (FE).

**Fig. S6.**
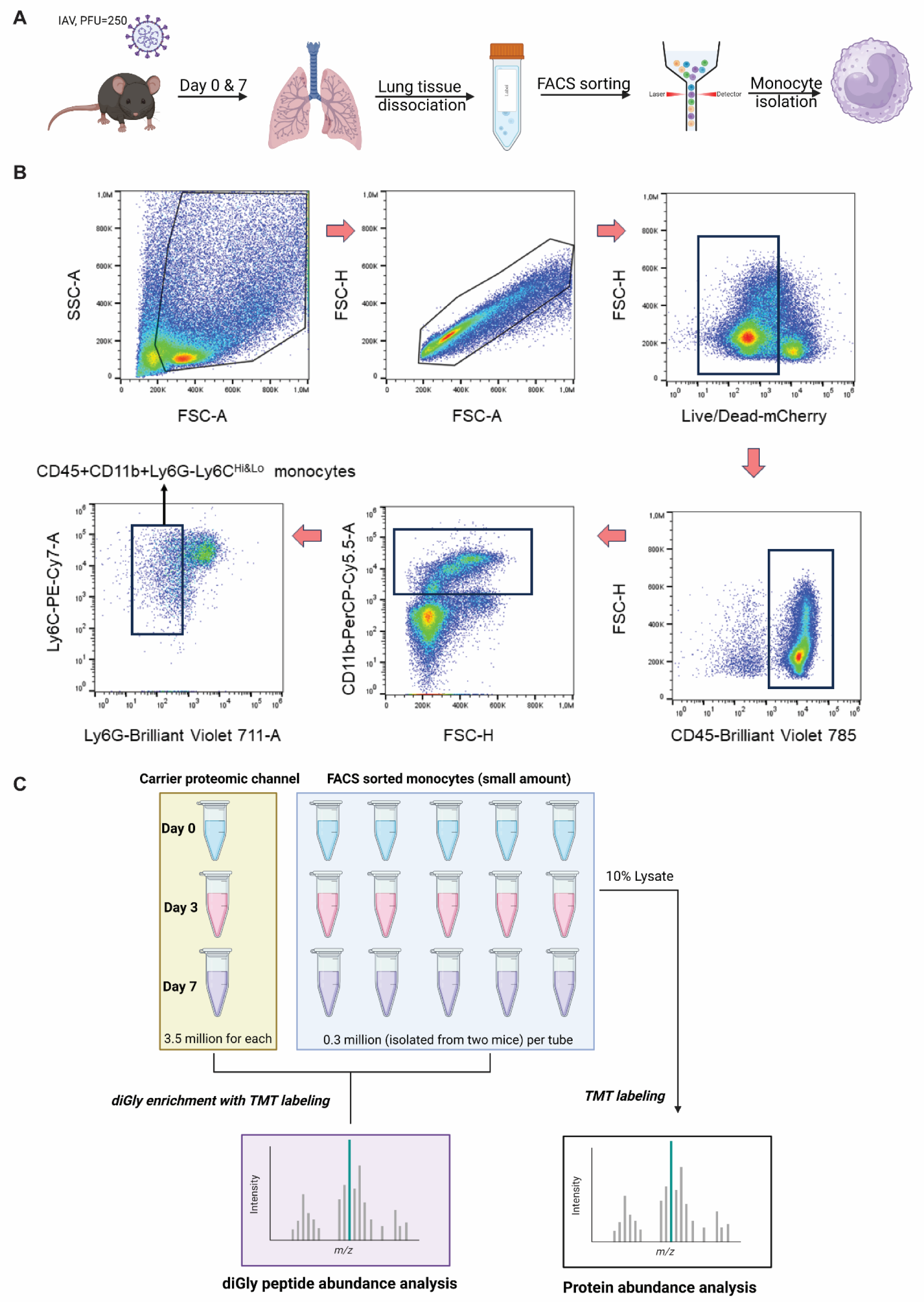
Experimental design for analysis of *in vivo* mouse monocytes. (**A**) Workflow for isolating monocytes from IAV infected mouse lungs. (**B**) FACS sorting strategy for isolating CD45+CD11b+Ly6G-Ly6C^Hi^ or CD45+CD11b+Ly6G-Ly6C^Lo^ monocytes from IAV infected mouse lungs. (**C**) Experimental design for the *in vivo* mouse monocytes’ proteomics analysis.

**Fig. S7.**
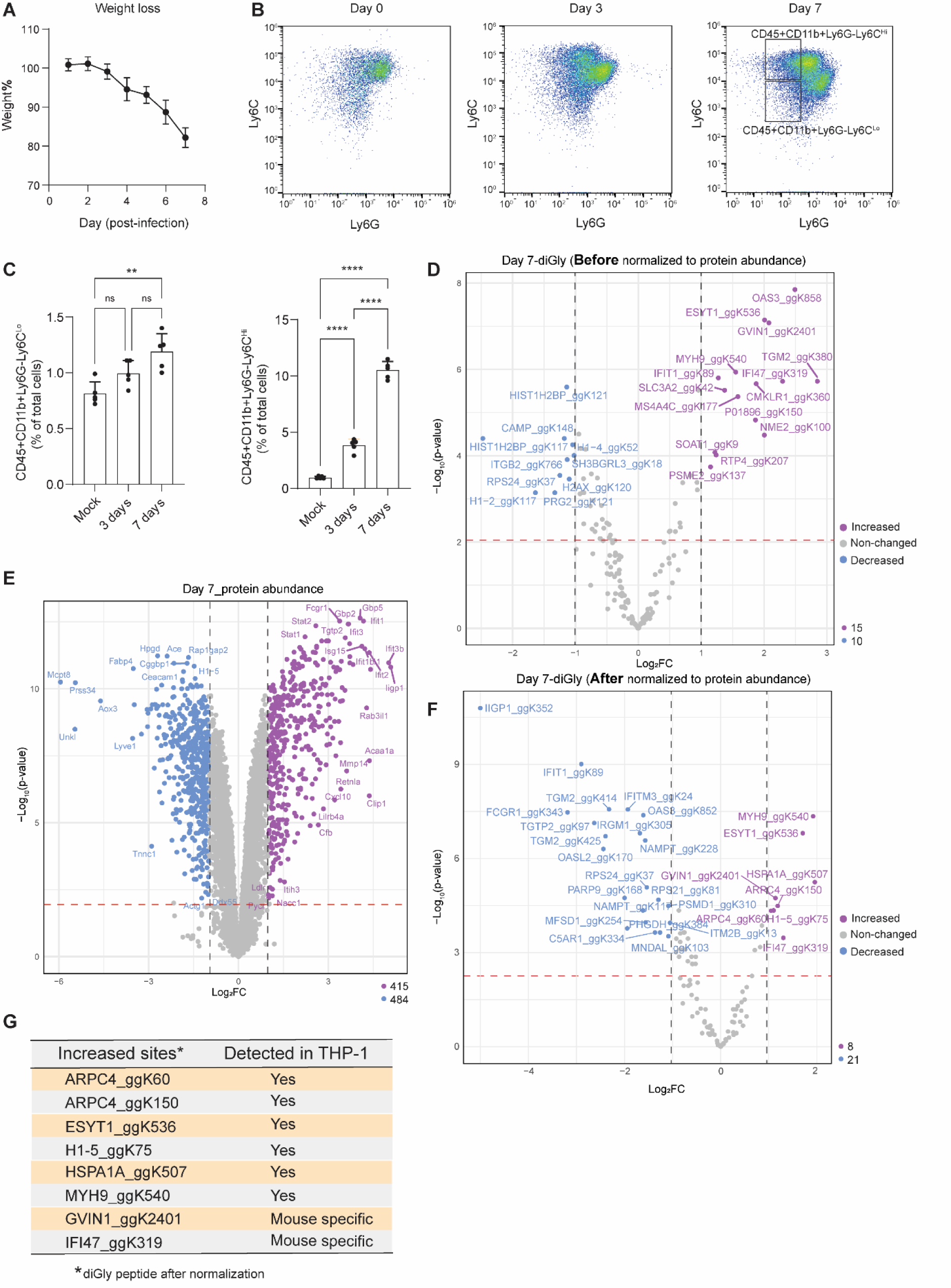
Ubiquitinated proteins in primary lung monocytes from IAV infected mice. (**A**) IAV (PR8, PFU = 250) induced mice weight loss (n = 10). Ratio was calculated as (measured weight/initial weight) × 100%. (**B**) Identification of CD45+CD11b+Ly6G-Ly6C^High^ or CD45+CD11b+Ly6G-Ly6C^Low^ monocytes in IAV infected mice lungs. (**C**) Quantification of CD45+CD11b+Ly6G-Ly6C^High^ or CD45+CD11b+Ly6G-Ly6C^Low^ monocytes from (B) (n = 10). Statistical analysis was performed with one-Way ANOVA test. **, p < 0.01; ****, p < 0.0001; ns, not significant. (**D**) Volcano plot of diGly peptides abundance before normalization to protein abundance. Dashed lines represent the log_2_FC = ±1 and p-value = 0.01. The number of increased or decreased proteins (log_2_FC > 1 or < -1, adj.p-value < 0.01) is indicated at the right bottom corner. (**E**) Volcano plot of total protein abundance variation induced by 7-day IAV infection in *in vivo* monocytes. Definitions of dashed lines and increased or decreased proteins are as in (D). (**F**) Volcano plot of diGly peptides abundance after normalization to protein abundance. Definitions of dashed lines and increased or decreased proteins are same as (D). (**G**) The induced diGly peptides (after normalization to protein abundance) with increased abundance (Log_2_FC > 1, adj.p-value < 0.01) in *in vivo* monocytes after IAV infection at Day 7 post-infection. Proteins and corresponding sites that were also identified in either Poly (I:C) treated (6 hr) or IAV infected (24 hr) THP-1 are marked “Yes” in “Detected in THP-1” column. “Mouse specific” means this protein is a unique gene in mice and lacks a human homolog.

**Fig. S8.**
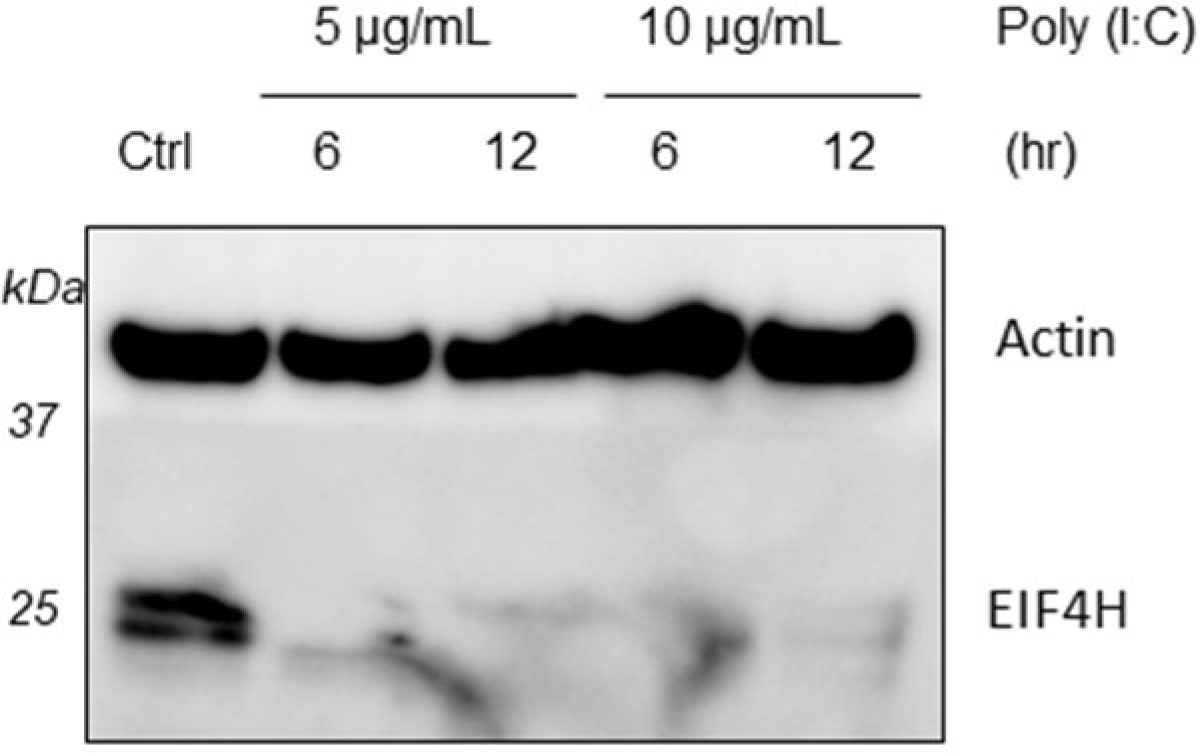
EIF4H degradation in A549 cells under Poly (I:C) treatment by immunoblotting. Poly (I:C) was transfected in A549 cells, and cells were collected at either 6 hr or 12 hr post transfection. Actin was used as an internal control. Ctrl: control, non-treated group (transfecting reagent, but no Poly (I:C) was added). Treatment conditions are indicated in the figure.

**Fig. S9.**
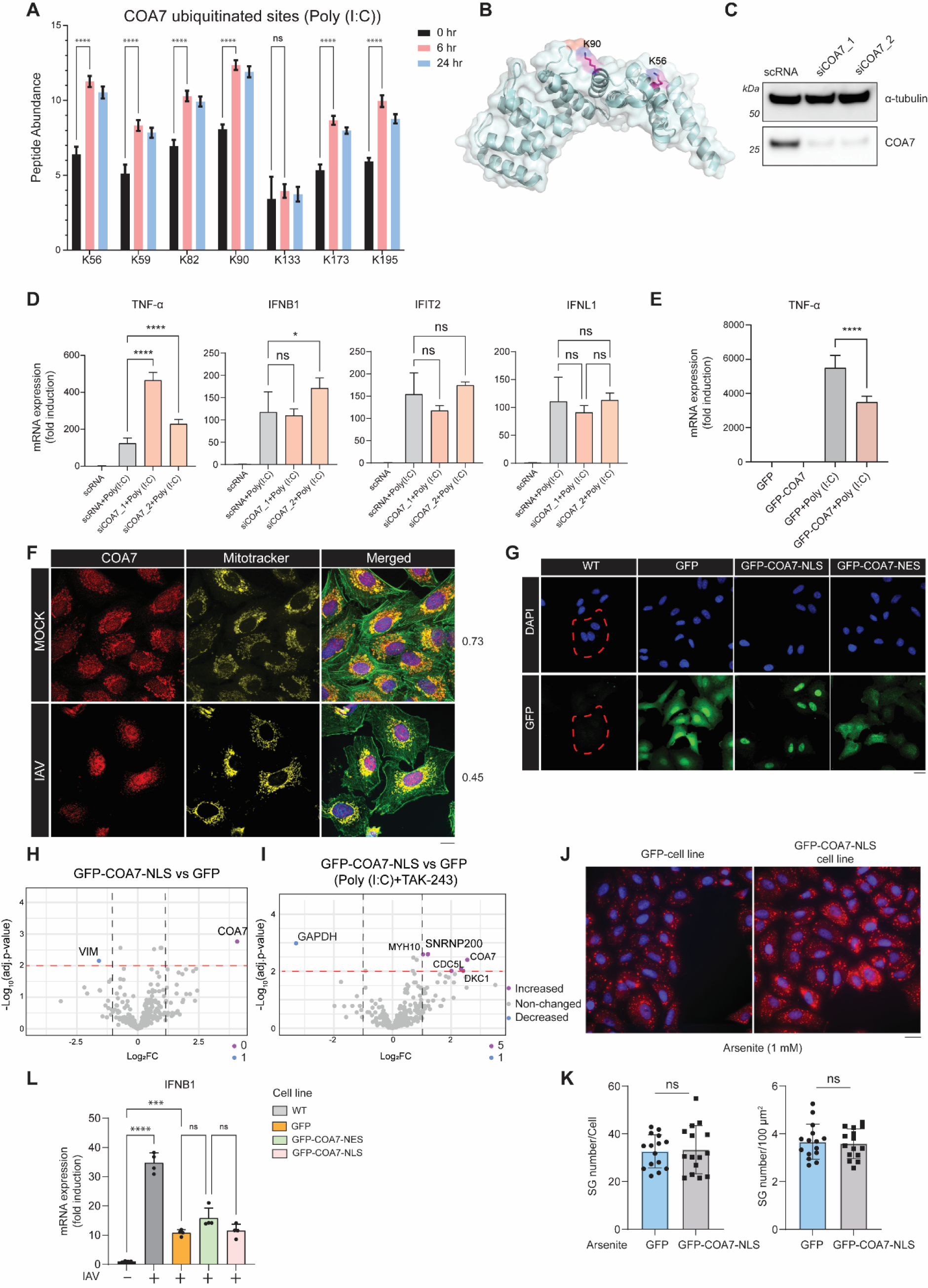
COA7 ubiquitination and its mechanism. (**A**) diGly peptides abundance of each COA7 lysine residue in Poly (I:C) treated (6 hr and 24 hr) THP-1 cells. Statistical analysis was performed with one-way ANOVA test. ****, p < 0.0001; ns, not significant. (**B**) COA7 K56 and K90 localization in the 3D structure. The COA7 structure was plotted by using PDB: 7MQZ. (**C**) COA7 siRNA knock-down in A549 cells. siRNAs were transfected in A549 WT cells with RNAiMAX, and cells were cultured for 48 hr before lysis and analyzed by immunoblotting. scRNA: scramble siRNA. α-tubulin was an internal control. (**D**) RT-qPCR quantification of TNF-α, IFNB1, IFIT2 and IFNL1 expression in COA7 knock-down A549 cells (n = 3). The 2^–ΔΔCt^ method was used for relative mRNA quantification. GAPDH was the reference gene. Statistical analysis was performed with one-way ANOVA test. *, p < 0.05; ****, p < 0.0001; ns, not significant. Poly (I:C) (5 μg/mL) was transfected in A549 cells with or without COA7 knock-down by siRNA (same as (C)). (**E**) RT-qPCR quantification of TNF-α expression in A549 cells stably overexpressing COA7 or GFP (n = 3). Methods and statistical analysis were as in (D). (**F**) Immunofluorescence of COA7 and mitochondria (visualized by Mito-Tracker) localization under IAV infection (X31, MOI = 1, 24 hr). Pearson’s R value (for evaluating the colocalization between COA7 signal and Mito-tracker signal) is indicated at the right. Scale bar is 20 µm. (**G**) Establishment of A549 cell lines expressing GFP, GFP-COA7-NLS and GFP-COA7-NES. Scale bar is 20 µm. Red dashed lines represent the cell boundary in WT A549 cells. (**H, I**) Volcano plot of GFP-COA7-NLS interacting proteins under non-stress condition (H) and Poly (I:C) +TAK-243 treatment (I). Treatment was same as in Figure 7. Dashed lines represent Log_2_FC = ±1, and adj.p-value = 0.01. The number of increased or decreased proteins (log_2_FC > 1 or < -1, adj.p-value < 0.01) are indicated at the right bottom corner (COA7 was excluded). (**J**) Immunofluorescence analysis of GFP-COA7-NLS effect on arsenite induced SGs formation. SGs (red) were visualized with G3BP1 antibody. Scale bar is 20 µm. (**K**) Quantification of SGs in (J). Statistical analysis was performed by unpaired t-test. ns, not significant. >10 pictures from 40x objective were used, and in total >400 cells for each group were quantified. (**L**) RT-qPCR of 24 hr IAV infection (X31, MOI = 1) induced IFNB1 expression in different A549 cell lines (n = 4). The 2^-ΔΔct^ method was used for relative mRNA quantification. GAPDH was used as reference gene. Statistical analysis was performed with one-way ANOVA test. *, p < 0.05; **, p < 0.01; ***, p < 0.001; ns, not significant.

**Table S1 to S10 (separate files)**

**Table S1. (separate file) TUBE label-free mass spectrometry.** Table showing results from TUBE proteomic analysis of THP-1 cells treated with Poly (I:C) and inhibitors listed in Figure 1D.

**Table S2. (separate file) TUBE-TMT mass spectrometry.** Table showing results from TUBE-TMT proteomic analysis of THP-1 cells treated with Poly (I:C) for different times listed in Figure 2A.

**Table S3. (separate file) diGly-TMT mass spectrometry for total protein abundance.** Table showing results from diGly-TMT proteomic analysis (only total protein abundance part) of THP-1 cells treated with Poly (I:C) for different times listed in Figure 3A.

**Table S4. (separate file) diGly-TMT Mass spectrometry for diGly peptides abundance.** Table showing results from diGly-TMT proteomic analysis (only diGly peptide part) of THP- 1 cells treated with Poly (I:C) for different times listed in Figure 3A.

**Table S5. (separate file) diGly-TMT mass spectrometry for IAV infected THP-1.** Table showing results from diGly-TMT proteomic analysis (both total protein abundance and diGly peptides parts) of IAV infected THP-1 cells for different times listed in Figure 4B.

**Table S6. (separate file) List of proteins with increased ubiquitination in IAV infected THP-1 cells at 24 hr.** Table showing results from diGly-TMT proteomic analysis (both total protein abundance and diGly peptides parts) of IAV infected THP-1 cells for different times listed in Figure 4G.

**Table S7. (separate file) diGly-TMT mass spectrometry for primary monocytes isolated from IAV infected mice at Day 7.** Table showing results from diGly-TMT proteomic analysis (both total protein abundance and diGly peptides parts) of primary monocytes CD45+CD11b+Ly6G-Ly6C^Hi^ in Figure S6C and S7D to F.

**Table S8. (separate file) Affinity purification-label free mass spectrometry for COA7 interacting proteins.** Table showing results from label free proteomic analysis of Poly (I:C) treated COA7 cell lines with or without TAK-243 treatment listed in Figure 7G to H and Figure S9 H to I.

**Table S9. (separate file) RNA sequencing qQUANT table.** RNA sequencing result for searching the gene expression listed in Figure S5.

**Table S10. (separate file) Primers for RT-qPCR and sequence of COA7 constructs.** Sequence details have been provided in the file.

## References and Notes

1. K. N. Swatek, D. Komander, Ubiquitin modifications. Cell Res 26, 399–422 (2016).

2. V. Chau et al., A multiubiquitin chain is confined to specific lysine in a targeted short-lived protein. Science 243, 1576–1583 (1989).

3. E. S. Johnson, P. C. Ma, I. M. Ota, A. Varshavsky, A proteolytic pathway that recognizes ubiquitin as a degradation signal. The Journal of biological chemistry 270, 17442–17456 (1995).

4. H. Hu, S. C. Sun, Ubiquitin signaling in immune responses. Cell Res 26, 457–483 (2016).

5. B. Gerlach et al., Linear ubiquitination prevents inflammation and regulates immune signalling. Nature 471, 591–596 (2011).

6. A. Werner et al., Cell-fate determination by ubiquitin-dependent regulation of translation. Nature 525, 523–527 (2015).

7. R. B. Damgaard, The ubiquitin system: from cell signalling to disease biology and new therapeutic opportunities. Cell Death Differ 28, 423–426 (2021).

8. N. Cusson-Hermance, S. Khurana, T. H. Lee, K. A. Fitzgerald, M. A. Kelliher, Rip1 mediates the Trif-dependent toll-like receptor 3- and 4-induced NF-{kappa}B activation but does not contribute to interferon regulatory factor 3 activation. The Journal of biological chemistry 280, 36560–36566 (2005).

9. P. N. Moynagh, The roles of Pellino E3 ubiquitin ligases in immunity. Nat Rev Immunol 14, 122–131 (2014).

10. H. Zhou et al., Bcl10 activates the NF-kappaB pathway through ubiquitination of NEMO. Nature 427, 167–171 (2004).

11. D. C. Scherer, J. A. Brockman, Z. Chen, T. Maniatis, D. W. Ballard, Signal-induced degradation of I kappa B alpha requires site-specific ubiquitination. Proc Natl Acad Sci U S A 92, 11259–11263 (1995).

12. T. Liu, L. Zhang, D. Joo, S. C. Sun, NF-κB signaling in inflammation. Signal Transduct Target Ther 2, 17023- (2017).

13. K. Pakos-Zebrucka et al., The integrated stress response. EMBO Rep 17, 1374–1395 (2016).

14. R. Jobava et al., Adaptive translational pausing is a hallmark of the cellular response to severe environmental stress. Molecular cell 81, 4191–4208.e4198 (2021).

15. R. Shalgi, J. A. Hurt, S. Lindquist, C. B. Burge, Widespread inhibition of posttranscriptional splicing shapes the cellular transcriptome following heat shock. Cell reports 7, 1362–1370 (2014).

16. K. Yamamoto et al., Control of the heat stress-induced alternative splicing of a subset of genes by hnRNP K. Genes Cells 21, 1006–1014 (2016).

17. V. Simões et al., Redox-sensitive E2 Rad6 controls cellular response to oxidative stress via K63-linked ubiquitination of ribosomes. Cell reports 39, 110860 (2022).

18. V. Balagopal, R. Parker, Polysomes, P bodies and stress granules: states and fates of eukaryotic mRNAs. Curr Opin Cell Biol 21, 403–408 (2009).

19. A. Buchberger, B. Bukau, T. Sommer, Protein quality control in the cytosol and the endoplasmic reticulum: brothers in arms. Molecular cell 40, 238–252 (2010).

20. B. A. Maxwell et al., Ubiquitination is essential for recovery of cellular activities after heat shock. Science 372, (2021).

21. H. Yang, H. Wang, J. Ren, Q. Chen, Z. J. Chen, cGAS is essential for cellular senescence. Proc Natl Acad Sci U S A 114, E4612–e4620 (2017).

22. M. Motwani, S. Pesiridis, K. A. Fitzgerald, DNA sensing by the cGAS-STING pathway in health and disease. Nat Rev Genet 20, 657–674 (2019).

23. J. Rehwinkel, M. U. Gack, RIG-I-like receptors: their regulation and roles in RNA sensing. Nat Rev Immunol 20, 537–551 (2020).

24. L. Alexopoulou, A. C. Holt, R. Medzhitov, R. A. Flavell, Recognition of double-stranded RNA and activation of NF-kappaB by Toll-like receptor 3. Nature 413, 732–738 (2001).

25. H. Kato et al., Differential roles of MDA5 and RIG-I helicases in the recognition of RNA viruses. Nature 441, 101–105 (2006).

26. WHO, World Health Organization 2023 data.who.int, WHO Coronavirus (COVID-19) dashboard > Cases [Dashboard]. https://data.who.int/dashboards/covid19/cases. ( 2023).

27. A. Stukalov et al., Multilevel proteomics reveals host perturbations by SARS-CoV-2 and SARS-CoV. Nature 594, 246–252 (2021).

28. G. Xu et al., Multiomics approach reveals the ubiquitination-specific processes hijacked by SARS-CoV-2. Signal Transduct Target Ther 7, 312 (2022).

29. R. T. Veenhuis et al., Monocyte-derived macrophages contain persistent latent HIV reservoirs. Nat Microbiol 8, 833–844 (2023).

30. J. Leon et al., A virus-specific monocyte inflammatory phenotype is induced by SARS-CoV-2 at the immune-epithelial interface. Proc Natl Acad Sci U S A 119, (2022).

31. E. Nikitina, I. Larionova, E. Choinzonov, J. Kzhyshkowska, Monocytes and Macrophages as Viral Targets and Reservoirs. Int J Mol Sci 19, (2018).

32. C. Junqueira et al., FcγR-mediated SARS-CoV-2 infection of monocytes activates inflammation. Nature 606, 576–584 (2022).

33. L. A. Perrone, J. K. Plowden, A. García-Sastre, J. M. Katz, T. M. Tumpey, H5N1 and 1918 pandemic influenza virus infection results in early and excessive infiltration of macrophages and neutrophils in the lungs of mice. PLoS Pathog 4, e1000115 (2008).

34. S. Vangeti et al., Human influenza virus infection elicits distinct patterns of monocyte and dendritic cell mobilization in blood and the nasopharynx. Elife 12, (2023).

35. T. Schmit et al., Interferon-gamma promotes monocyte-mediated lung injury during influenza infection. Cell reports 38, 110456 (2022).

36. B. M. Coates et al., Inflammatory Monocytes Drive Influenza A Virus-Mediated Lung Injury in Juvenile Mice. J Immunol 200, 2391–2404 (2018).

37. The new scope of virus taxonomy: partitioning the virosphere into 15 hierarchical ranks. Nat Microbiol 5, 668-674 (2020).

38. Y. Yoshida et al., A comprehensive method for detecting ubiquitinated substrates using TR-TUBE. Proc Natl Acad Sci U S A 112, 4630–4635 (2015).

39. H. Tsuchiya et al., Ub-ProT reveals global length and composition of protein ubiquitylation in cells. Nature communications 9, 524 (2018).

40. A. Shaalan, G. Carpenter, G. Proctor, Caspases are key regulators of inflammatory and innate immune responses mediated by TLR3 in vivo. Mol Immunol 94, 190–199 (2018).

41. R. Qi, D. Singh, C. C. Kao, Proteolytic processing regulates Toll-like receptor 3 stability and endosomal localization. The Journal of biological chemistry 287, 32617–32629 (2012).

42. Y. Wang et al., Inflammasome Activation Triggers Caspase-1-Mediated Cleavage of cGAS to Regulate Responses to DNA Virus Infection. Immunity 46, 393–404 (2017).

43. I. Tapescu et al., The RNA helicase DDX39A binds a conserved structure in chikungunya virus RNA to control infection. Molecular cell 83, 4174–4189.e4177 (2023).

44. S. X. Ge, D. Jung, R. Yao, ShinyGO: a graphical gene-set enrichment tool for animals and plants. Bioinformatics 36, 2628–2629 (2020).

45. W. Kim et al., Systematic and quantitative assessment of the ubiquitin-modified proteome. Molecular cell 44, 325–340 (2011).

46. J. Peng et al., A proteomics approach to understanding protein ubiquitination. Nat Biotechnol 21, 921–926 (2003).

47. G. Xu, J. S. Paige, S. R. Jaffrey, Global analysis of lysine ubiquitination by ubiquitin remnant immunoaffinity profiling. Nat Biotechnol 28, 868–873 (2010).

48. F. Li, et al., Monocyte-derived alveolar macrophages autonomously determine severe outcome of respiratory viral infection. Sci Immunol 7, eabj5761 (2022).

49. A. A. Petelski et al., Multiplexed single-cell proteomics using SCoPE2. Nat Protoc 16, 5398–5425 (2021).

50. G. T. Ellis et al., TRAIL+ monocytes and monocyte-related cells cause lung damage and thereby increase susceptibility to influenza-Streptococcus pneumoniae coinfection. EMBO Rep 16, 1203–1218 (2015).

51. Z. A. Jaafar, J. S. Kieft, Viral RNA structure-based strategies to manipulate translation. Nat Rev Microbiol 17, 110–123 (2019).

52. D. Flierman, C. S. Coleman, C. M. Pickart, T. A. Rapoport, V. Chau, E2-25K mediates US11-triggered retro-translocation of MHC class I heavy chains in a permeabilized cell system. Proc Natl Acad Sci U S A 103, 11589–11594 (2006).

53. M. Elangovan, H. K. Chong, J. H. Park, E. J. Yeo, Y. J. Yoo, The role of ubiquitin-conjugating enzyme Ube2j1 phosphorylation and its degradation by proteasome during endoplasmic stress recovery. J Cell Commun Signal 11, 265–273 (2017).

54. L. E. Formosa et al., Mitochondrial COA7 is a heme-binding protein with disulfide reductase activity, which acts in the early stages of complex IV assembly. Proc Natl Acad Sci U S A 119, (2022).

55. E. K. A. Nur et al., Nuclear translocation of cytochrome c during apoptosis. The Journal of biological chemistry 279, 24911–24914 (2004).

56. R. M. Kluck, E. Bossy-Wetzel, D. R. Green, D. D. Newmeyer, The release of cytochrome c from mitochondria: a primary site for Bcl-2 regulation of apoptosis. Science 275, 1132–1136 (1997).

57. M. Paget et al., Stress granules are shock absorbers that prevent excessive innate immune responses to dsRNA. Molecular cell 83, 1180–1196.e1188 (2023).

58. L. Wang et al., Disrupting the HDAC6-ubiquitin interaction impairs infection by influenza and Zika virus and cellular stress pathways. Cell reports 39, 110736 (2022).

59. D. S. W. Protter, R. Parker, Principles and Properties of Stress Granules. Trends Cell Biol 26, 668–679 (2016).

60. Y. Liao et al., UBAP2L ensures homeostasis of nuclear pore complexes at the intact nuclear envelope. The Journal of cell biology 223, (2024).

61. T. Ikeda, K. Yamazaki, F. Okumura, T. Kamura, K. Nakatsukasa, Role of the San1 ubiquitin ligase in the heat stress-induced degradation of nonnative Nup1 in the nuclear pore complex. Genetics 226, (2024).

62. C. Boyault et al., HDAC6 controls major cell response pathways to cytotoxic accumulation of protein aggregates. Genes & development 21, 2172–2181 (2007).

63. S. Hirayama et al., Nuclear export of ubiquitinated proteins via the UBIN-POST system. Proc Natl Acad Sci U S A 115, E4199–e4208 (2018).

64. Y. Ogawa, N. Imamoto, Nuclear transport adapts to varying heat stress in a multistep mechanism. The Journal of cell biology 217, 2341–2352 (2018).

65. K. Zhang et al., Stress Granule Assembly Disrupts Nucleocytoplasmic Transport. Cell 173, 958–971 e917 (2018).

66. M. Mattern, J. Sutherland, K. Kadimisetty, R. Barrio, M. S. Rodriguez, Using Ubiquitin Binders to Decipher the Ubiquitin Code. Trends Biochem Sci 44, 599–615 (2019).

67. D. Komander, M. Rape, The ubiquitin code. Annu Rev Biochem 81, 203–229 (2012).

68. C. Hoege, B. Pfander, G. L. Moldovan, G. Pyrowolakis, S. Jentsch, RAD6-dependent DNA repair is linked to modification of PCNA by ubiquitin and SUMO. Nature 419, 135–141 (2002).

69. B. D. Freudenthal, L. Gakhar, S. Ramaswamy, M. T. Washington, Structure of monoubiquitinated PCNA and implications for translesion synthesis and DNA polymerase exchange. Nat Struct Mol Biol 17, 479–484 (2010).

70. M. Bienko et al., Ubiquitin-binding domains in Y-family polymerases regulate translesion synthesis. Science 310, 1821–1824 (2005).

71. M. Bienko et al., Regulation of translesion synthesis DNA polymerase eta by monoubiquitination. Molecular cell 37, 396–407 (2010).

72. J. Terrell, S. Shih, R. Dunn, L. Hicke, A function for monoubiquitination in the internalization of a G protein-coupled receptor. Molecular cell 1, 193–202 (1998).

73. D. K. Stringer, R. C. Piper, A single ubiquitin is sufficient for cargo protein entry into MVBs in the absence of ESCRT ubiquitination. The Journal of cell biology 192, 229–242 (2011).

74. M. Li et al., Mono-versus polyubiquitination: differential control of p53 fate by Mdm2. Science 302, 1972–1975 (2003).

75. J. Yuan, K. Luo, L. Zhang, J. C. Cheville, Z. Lou, USP10 regulates p53 localization and stability by deubiquitinating p53. Cell 140, 384–396 (2010).

76. N. Zheng, N. Shabek, Ubiquitin Ligases: Structure, Function, and Regulation. Annu Rev Biochem 86, 129–157 (2017).

77. D. Oudshoorn et al., HERC6 is the main E3 ligase for global ISG15 conjugation in mouse cells. PloS one 7, e29870 (2012).

## References and Notes

1. H. Tsuchiya et al., Ub-ProT reveals global length and composition of protein ubiquitylation in cells. Nature communications 9, 524 (2018).

2. S. Tyanova, T. Temu, J. Cox, The MaxQuant computational platform for mass spectrometry-based shotgun proteomics. Nat Protoc 11, 2301–2319 (2016).

3. T. Welte et al., Convergence of multiple RNA-silencing pathways on GW182/TNRC6. Molecular cell 83, 2478–2492.e2478 (2023).

4. Y. Wang et al., Reversed-phase chromatography with multiple fraction concatenation strategy for proteome profiling of human MCF10A cells. Proteomics 11, 2019–2026 (2011).

5. The Galaxy platform for accessible, reproducible, and collaborative data analyses: 2024 update. Nucleic Acids Res 52, W83-w94 (2024).

6. D. Gaidatzis, A. Lerch, F. Hahne, M. B. Stadler, QuasR: quantification and annotation of short reads in R. Bioinformatics 31, 1130–1132 (2015).

7. E. Campeau et al., A versatile viral system for expression and depletion of proteins in mammalian cells. PloS one 4, e6529 (2009).

